# The actin nucleation factors JMY and WHAMM enable a rapid p53-dependent pathway of apoptosis

**DOI:** 10.1101/2020.07.15.205518

**Authors:** Virginia L. King, Nathan K. Leclair, Kenneth G. Campellone

## Abstract

The actin cytoskeleton is a well-known player in most vital cellular processes, but comparably little is understood about how the actin assembly machinery impacts programmed cell death pathways. In the current study, we explored roles for the human <u>W</u>iskott-<u>A</u>ldrich <u>S</u>yndrome <u>P</u>rotein (WASP) family of actin nucleation factors in DNA damage-induced apoptosis. Inactivation of each WASP-family gene revealed that two, *JMY* and *WHAMM*, are required for rapid apoptotic responses. JMY and WHAMM enable p53-dependent cell death by enhancing mitochondrial permeabilization, initiator caspase cleavage, and executioner caspase activation. The loss of JMY additionally results in significant changes in gene expression, including upregulation of the small G-protein RhoD. Depletion or deletion of *RHOD* increases cell death, suggesting that RhoD normally plays a key role in cell survival. These results give rise to a model in which JMY and WHAMM promote intrinsic cell death responses that can be opposed by RhoD.

**Author Summary:** The actin cytoskeleton is a collection of protein polymers that assemble and disassemble within cells at specific times and locations. Cytoskeletal regulators called nucleation-promoting factors ensure that actin polymerizes when and where it is needed, and many of these factors are members of the <u>W</u>iskott-<u>A</u>ldrich <u>S</u>yndrome <u>P</u>rotein (WASP) family. Humans express 8 WASP-family proteins, but whether the different factors function in programmed cell death pathways is not well understood. In this study, we explored roles for each WASP-family member in apoptosis and found that a subfamily consisting of JMY and WHAMM are critical for a rapid pathway of cell death. Furthermore, the loss of JMY results in changes in gene expression, including a dramatic upregulation of the small G-protein RhoD, which appears to be crucial for cell survival. Collectively, our results point to the importance of JMY and WHAMM in driving intrinsic cell death responses plus a distinct function for RhoD in maintaining cell viability.

## Introduction

Apoptosis is a programmed form of cell death crucial for many organismal processes, including development, tissue turnover, and tumor suppression [1, 2]. It is driven by intrinsic mitochondria-mediated and extrinsic receptor-mediated death pathways that converge on a terminal execution program [3–6]. Cell rounding, shrinkage, membrane blebbing, and fragmentation into apoptotic bodies are common morphological features of apoptosis, and are controlled by a loss of actin-associated adhesions, rearrangements of actin filaments, and actin depolymerization [7, 8]. While this reorganization and disassembly of the cytoskeleton during apoptosis is well described, the extent to which the actin assembly machinery contributes to the initiation or progression of apoptotic pathways is not understood.

Actin polymerization into filaments has been extensively characterized during cell morphogenesis, intracellular trafficking, cytokinesis, and motility [9, 10]. To assemble branched actin networks during these processes, the heptameric Arp2/3 complex cooperates with nucleation-promoting factors from the Wiskott-Aldrich Syndrome Protein (WASP) family [11, 12]. The mammalian WASP family is composed of WASP, N-WASP, WAVE1, WAVE2, WAVE3, WASH, WHAMM, JMY, and WHIMP [13, 14]. Several of these factors are found in multi-protein complexes themselves, including all three WAVE isoforms and WASH, which constitute the WAVE and WASH complexes [15, 16]. Each WASP-family member uses a conserved C-terminal domain that binds actin monomers and the Arp2/3 complex to stimulate polymerization, while their divergent N-terminal domains interact with a variety of small G-proteins and phospholipids which direct their different localizations and functions within cells [9, 13]. N-WASP, WAVE1-3, and WHIMP drive plasma membrane protrusion; N-WASP and WASH control endocytosis and endosomal cargo sorting; WHAMM and JMY promote anterograde membrane transport and autophagy. Additionally, during most of these processes, the atypical nucleation-promoting factor Cortactin binds actin filaments and the Arp2/3 complex to modulate actin branchpoint stability [17].

Compared to their well-described roles in such vital cellular functions, the extent to which WASP-family proteins participate in cell death pathways is largely uncharacterized. WAVE1 is perhaps the most studied member in relation to apoptosis, where it can influence the localization or modification of Bcl-2-family proteins, which control mitochondrial outer membrane permeabilization and the release of apoptogenic proteins [6, 18, 19]. In hepatocytes, WAVE1 forms a complex with the pro-apoptotic protein Bad [20], while in neuronal cells WAVE1 interacts with the anti-apoptotic protein Bcl-xL to promote mitochondrial recruitment of the pro-apoptotic permeabilization factor Bax [21]. In contrast, in leukemia cells, WAVE1 interacts with the anti-apoptotic protein Bcl-2 such that WAVE1 overexpression inhibits death signaling, whereas WAVE1 depletion increases apoptosis in response to chemotherapeutic agents [22, 23]. Thus, WAVE1 appears to have cell type specific effects on apoptotic processes.

In addition to WAVE1, JMY has been reported to function in apoptosis. JMY was discovered as a cofactor that modulates the function of p53 [24], a central tumor suppressor protein and transcription factor [25, 26]. Under normal cellular growth conditions, JMY is maintained in the cytoplasm through interactions with the ubiquitin ligase Mdm2 [27] as well as autophagy-related proteins [28]. Upon genotoxic damage, JMY binds importins and accumulates in the nucleus [29, 30], where it associates with the stress-response protein Strap, the histone acetyltransferase p300, and p53 [24, 31]. Co-overexpression of JMY and p300 in p53-proficient epithelial cells drives increased transcription of Bax [24, 29], while introduction of a JMY siRNA in p53-overexpressing cells decreases the amount of Bax induction [29]. These data indicate that JMY can function in apoptosis by enhancing the transcription of at least one pro-apoptotic gene. Consistent with this conclusion, an siRNA targeting JMY causes a decrease in the percentage of sub-G1 (presumably apoptotic) cells after UV treatment [27]. Although, under other conditions, siRNA targeting of JMY can increase the number of sub-G1 cells [32].

Collectively, these studies on WAVE1 and JMY reveal an interesting dynamic between actin assembly proteins and apoptosis-associated processes, and suggest that the relationships between nucleation factors and cell death pathways merit further investigation. In the current study, using a systematic gene knockout approach, we tested roles for all WASP-family members in DNA damage-induced apoptosis. We discovered that JMY and WHAMM are key regulators of a rapid p53-dependent cell death pathway, but that their pro-apoptotic functions are opposed by the small G-protein RhoD.

## Results

### The WASP-family proteins JMY and WHAMM are important for apoptosis

To evaluate the impact of WASP-family members on apoptosis, we employed a panel of knockout (KO) HAP1 or eHAP fibroblast-like cell lines lacking N-WASP, WAVE1, WAVE2, WAVE3, the WAVE Complex, the WASH Complex, WHAMM, JMY, or Cortactin (S1 Table). These cells, derived from a chronic myeloid leukemia patient [33], are useful for studying apoptosis because they harbor an immortalizing BCR-ABL fusion [34], have easily manipulatable nearly-haploid (HAP1) or fully haploid (eHAP) genomes [33, 35], and are p53-proficient [36]. To examine intrinsic apoptotic responses, the parental and KO cell lines were treated with etoposide, a topoisomerase II inhibitor that induces double-strand DNA breaks. To monitor cell death, each line was then imaged using fluorescent annexin V (AnnV) to identify the apoptotic hallmark of phosphatidylserine externalization, propidium iodide (PI) to assess membrane permeability, and hoescht to visualize nuclear fragmentation (Fig 1A; S1 Fig; S1 Video).

**Fig 1.**
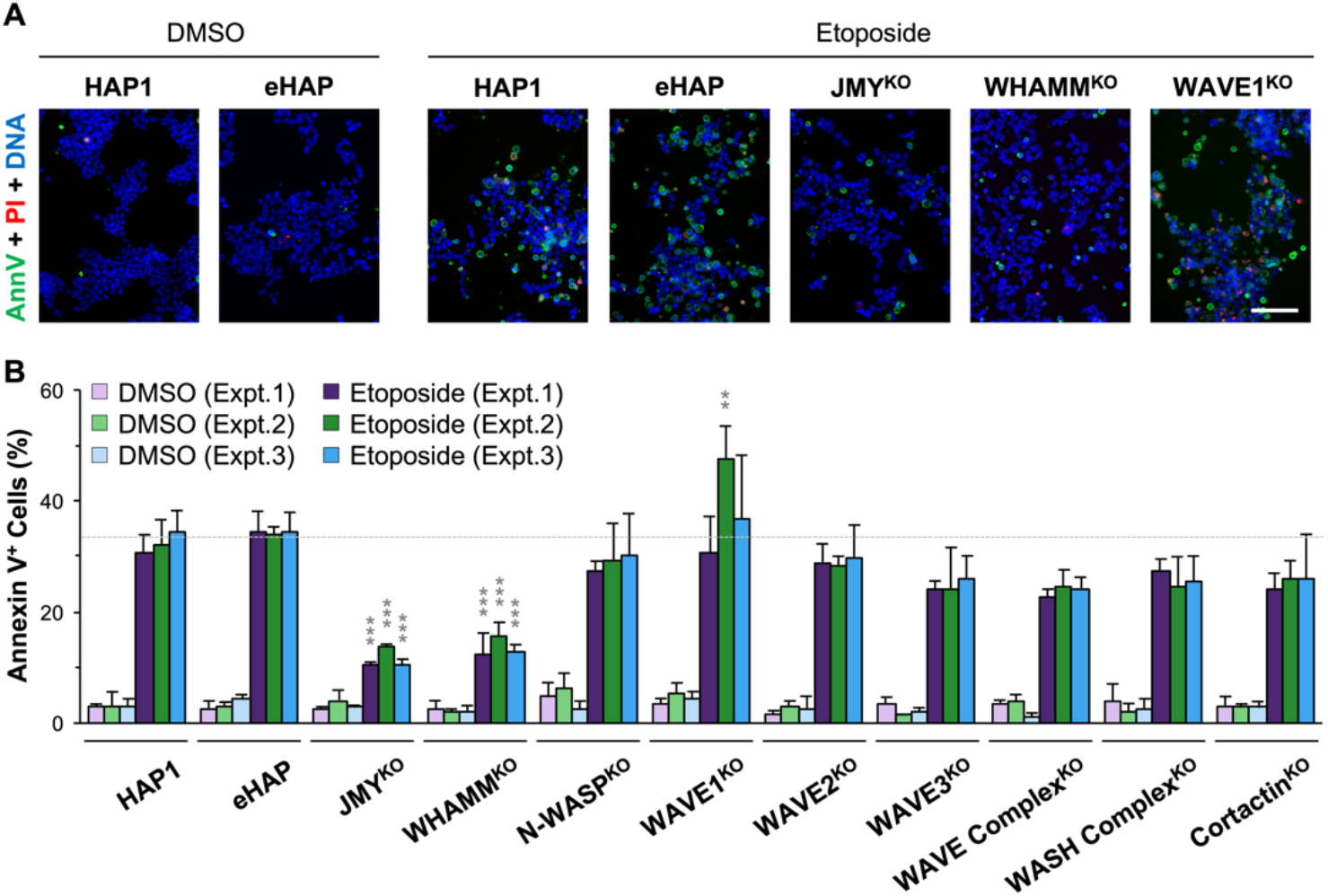
Cells lacking the WASP-family members JMY or WHAMM undergo less apoptosis. **(A)** Parental (HAP1, eHAP) and WASP-family knockout (JMY^KO^, WHAMM^KO^, N-WASP^KO^, WAVE1^KO^, WAVE2^KO^, WAVE3^KO^, WAVE Complex^KO^, WASH Complex^KO^, Cortactin^KO^) cells were treated with DMSO or 5μM etoposide for 6h and stained with Alexa488-AnnexinV (AnnV; green), Propidium Iodide (PI; red), and Hoescht (DNA; blue). Scale bar, 100μm. **(B)** The % of AnnV-positive cells was calculated in ImageJ by dividing the number of cells that displayed AnnV staining by the total number of cells identified by nuclear Hoescht staining. Each bar represents the mean ±SD from 3 fields-of-view in 3 separate experiments (n = 414-1,303 cells per bar). The gray dashed line depicts the mean %AnnV-positive cells for the etoposide-treated HAP1 and eHAP parental lines. Significance stars are for comparisons to the etoposide-treated HAP1 or eHAP cells. **p<0.01, ***p<0.001 (ANOVA, Tukey post-hoc tests).

After 6h of etoposide treatment, approximately 33% of parental HAP1 or eHAP cells were apoptotic, as determined by AnnV-positive staining, compared to only 3% of DMSO-treated controls (Fig 1B). The proportion of cells displaying AnnV staining was not significantly different across the majority of the etoposide-or DMSO-treated WASP-family knockout lines compared to the parental lines (Fig 1B). However in one of three etoposide experiments, a statistically significant increase in AnnV-positive cells was observed for the WAVE1^KO^ line (Fig 1B). This phenotype is reminiscent of previous studies in which WAVE1 depletion in leukemia cells increased apoptosis [22, 23]. In contrast, two cell lines, the JMY^KO^ and the WHAMM^KO^, exhibited significantly lower percentages of AnnV-positive cells across all three etoposide experiments (Fig 1A and 1B). The JMY^KO^ and WHAMM^KO^ cell lines also showed substantially less frequent PI staining and nuclear fragmentation compared to parental cells (Fig 1A; S1 Fig). These results indicate that among a panel of cell lines lacking WASP-family members, only those missing JMY or WHAMM have significantly impaired apoptotic responses to DNA damage.

### JMY and WHAMM enable DNA damage-induced apoptosis in several cellular contexts

JMY and WHAMM are approximately 35% identical and 50% similar to one another [37], and comprise a subgroup within the WASP family. Since both appear to be important for apoptosis, we sought to more thoroughly compare the effects of *JMY* or *WHAMM* inactivation on cell death. For this, we characterized three independent *JMY* mutant cell lines derived from HAP1 cells and two independent *WHAMM* mutant cell lines derived from eHAP cells (Fig 2A and 2B; S2 Fig). Etoposide treatment caused similarly high levels of DNA damage across all of the parental and knockout cell lines, as evidenced by increased phosphorylated histone H2AX (γH2AX) protein levels and greater numbers of γH2AX foci at DNA breaks within nuclei (S3 Fig). Both the parental cell lines, as well as the JMY^KO^ and the WHAMM^KO^ lines, showed etoposide dose-dependent increases in AnnV-positive apoptotic cells, but the mutants displayed significantly lower percentages at virtually every concentration (Fig 2C and 2D). At the highest concentrations (4-5μM), nearly 40% of parental cells, but only 10-15% of JMY^KO^ cells and 15-20% of WHAMM^KO^ cells, were AnnV-positive (Fig 2C and 2D).

**Fig 2.**
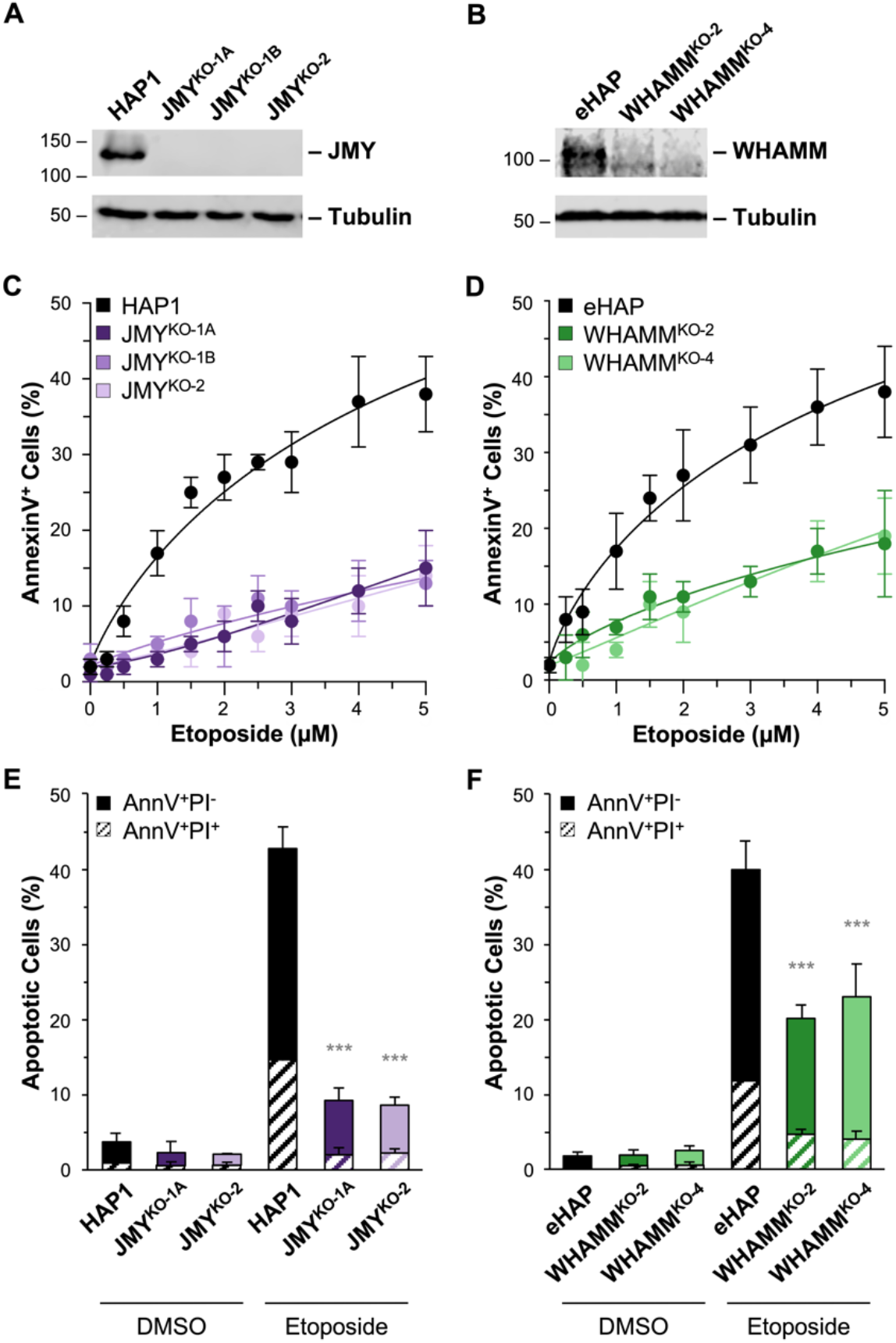
JMY- and WHAMM-deficient cells have defective apoptotic responses following DNA damage. **(A-B)** HAP1, JMY^KO^, eHAP, and WHAMM^KO^ cell lysates were immunoblotted with anti-JMY, anti-WHAMM, and anti-tubulin antibodies. **(C-D)** HAP1, JMY^KO^, eHAP, and WHAMM^KO^ cells were treated with a range of etoposide concentrations for 6h and then stained with Alexa488-AnnV and Hoescht. The % of AnnV-positive cells was calculated in ImageJ. Each point represents the mean ±SD from 3-6 fields-of-view pooled from 1-2 experiments (n = 1,699-4,276 cells per experiment in C; n = 2,896-3,572 cells per experiment in D). Nonlinear regressions were performed with a baseline set to 0.02 and a maximum response set to 0.85. The EC50 is significantly different for parental vs KO samples (EC50: Parental = 6.1-6.4 μM; JMY^KO^ = 16.6-39.7 μM, p<0.001; WHAMM^KO^ = 16.2-32.7 μM, p<0.001). **(E-F)** Cells were treated with DMSO or 5μM etoposide for 6h and stained with Alexa488-AnnV, PI, and Hoescht. The % of AnnV-positive cells was calculated and displayed as the fraction of AnnV-positive/PI-negative (AnnV^+^PI^−^) or AnnV/PI double-positive (AnnV^+^PI^+^) cells. Each bar represents the mean ±SD from 3 experiments (n = 5,206-8,876 cells per bar in E; n = 2,538-4,878 cells per bar in F). Significance stars refer to comparisons of total AnnV^+^ counts for parental vs KO samples. ***p<0.001 (ANOVA, Tukey post-hoc tests).

To evaluate early versus late apoptosis, the percentage of AnnV-positive cells without versus with PI staining was quantified (S1 Fig). Among the ~40% of parental cells exhibiting AnnV staining, about 25% were AnnV-positive and PI-negative, indicating early apoptosis, while 15% were AnnV and PI double-positive, signifying late apoptosis (Fig 2E and 2F). In comparison, all of the JMY^KO^ and WHAMM^KO^ samples contained significantly fewer cells in both early and late apoptosis, with the JMY^KO^ cells exhibiting 8% early and 2% late (Fig 2E), and the WHAMM^KO^ cells displaying 15% early and 5% late (Fig 2F). These results indicate that JMY and WHAMM each play important roles in apoptosis, but the more extreme phenotypes observed when *JMY* is mutated suggest that JMY is more prominent in enabling cell death.

To determine if transient depletion of JMY also inhibits apoptosis, we next treated parental HAP1 cells with two independent siRNAs targeting JMY (Fig 3A). After etoposide exposure, significantly fewer JMY-depleted cells were AnnV-positive compared to control cells (Fig 3B and 3C), and the amount of JMY expressed in the cells positively correlated with the percentage of AnnV-positive cells (Fig 3D). To additionally gauge apoptotic phenotypes in different human cell lines, we targeted JMY using siRNAs in both U2OS osteosarcoma cells and HeLa adenocarcinoma cells. Again, after etoposide exposure, depletion of JMY resulted in significantly fewer AnnV-positive cells compared to control siRNA treatments (S4 Fig). Moreover, the level of JMY expressed in U2OS and HeLa cells showed a positive relationship with the proportion of AnnV-positive cells (S4 Fig). These results demonstrate that JMY is required for efficient apoptosis across multiple cell types.

**Fig 3.**
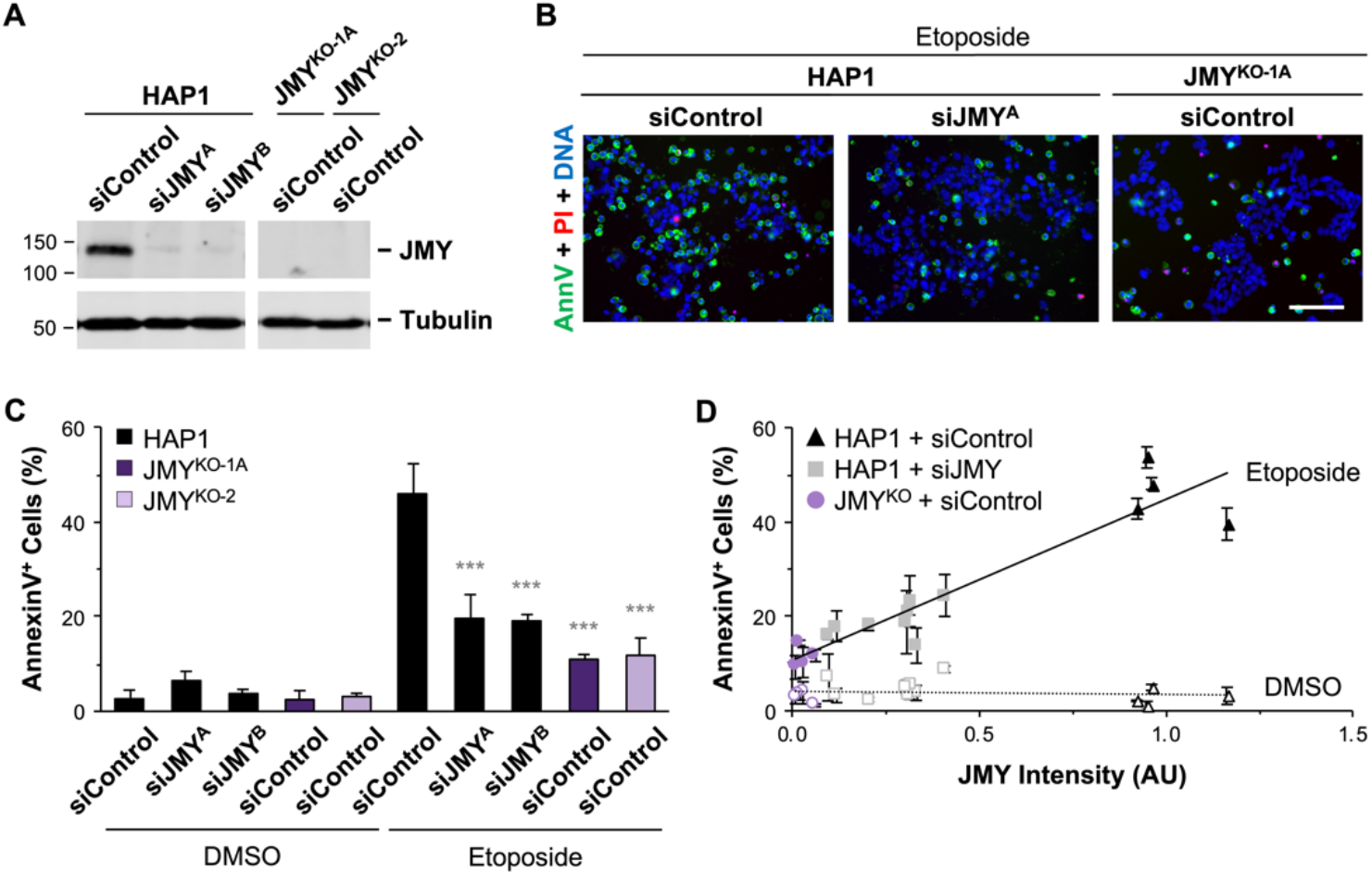
Transient depletion of JMY results in less apoptosis in response to DNA damage. **(A)** HAP1 cells were treated with control siRNAs or independent siRNAs for the JMY gene, while JMY^KO^ cell lines were treated with control siRNAs before immunoblotting with anti-JMY and anti-tubulin antibodies. **(B)** Cells were treated with DMSO or 5μM etoposide for 6h and stained with Alexa488-AnnV (green), PI (red), and Hoescht (blue). Scale bar, 100μm. **(C)** The % of AnnV-positive cells was calculated and each bar represents the mean ±SD from 2-4 experiments (n = 2,789-6,207 cells per bar). Significance stars refer to comparisons to the HAP1 siControl sample. **(D)** JMY band intensities on immunoblots were normalized to tubulin bands and plotted against the % of AnnV-positive cells. Each point represents the mean ±SD from 3 fields-of-view in a given experiment (n = 564-2,387 cells per point). The linear trendline regression equation for etoposide-treated samples (Y = 34.56X + 10.52) was significantly non-zero (p<0.001, R^2^=0.86). ***p<0.001 (ANOVA, Tukey post-hoc tests).

### Inactivation of *JMY* or *WHAMM* inhibits multiple steps of intrinsic apoptotic signaling

Since the loss of JMY or WHAMM decreased the final apoptotic readouts of phosphatidylserine externalization and membrane permeability, we next wanted to determine if the inactivation of *JMY* or *WHAMM* affected earlier aspects of apoptosis. Following acute DNA damage, apoptotic signaling cascades are typified by the multimerization and proteolytic cleavage of initiator caspases that cleave and activate executioner caspases, which in turn target multiple proteins in the cytosol and nucleus [38, 39]. Terminal executioner caspase-3 and caspase-7 cleave the motif Asp-Glu-Val-Asp (DEVD) [39], so we used a DEVD substrate peptide conjugated to a fluorescent reporter to quantify the amount of cells with active caspase-3/7 following treatment with etoposide for 3h or 6h. Caspase-3/7 activation was readily detectable in parental cells at 3h, but did not become clearly apparent in the knockout cells until the 6h timepoint (Fig 4A and 4B), when 50% of parental cells versus only 15-25% of JMY^KO^ or WHAMM^KO^ cells were positive for cleaved DEVD (Fig 4C and 4D). Therefore, the apoptotic defects arising from *JMY* or *WHAMM* mutations include significantly delayed and less potent activation of executioner caspases.

**Fig 4.**
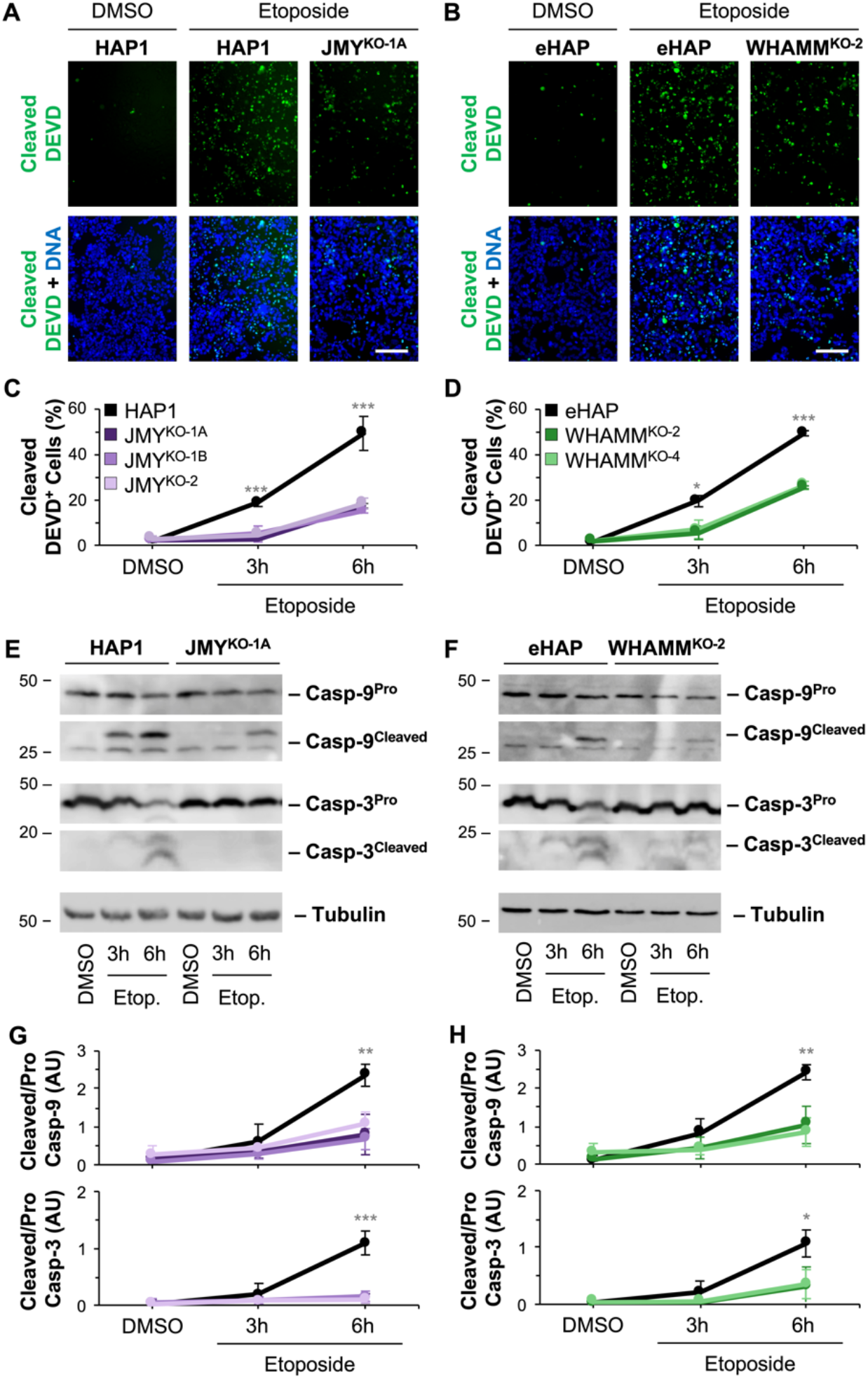
Initiator and executioner caspase cleavage is inefficient in the absence of JMY or WHAMM. **(A-B)** HAP1, JMY^KO^, eHAP, and WHAMM^KO^ cells were treated with DMSO for 6h or 5μM etoposide for 3h or 6h and stained with caspase-3/7 green detection reagent to label cleaved DEVD (green) and Hoescht to stain DNA (blue). Images are from the 6h timepoint. Scale bars, 100μm. **(C-D)** The % of cleaved DEVD-positive cells was calculated in ImageJ by counting cells that exhibited green nuclear fluorescence and dividing by the total number of Hoescht-stained nuclei. Each point represents the mean ±SD from 3 experiments (n = 3,063-6,100 cells per point in C; n = 2,741-6,646 cells per point in D). Significance stars refer to comparisons of parental to KO samples at the depicted timepoints. **(E-F)** Cells were treated with DMSO or etoposide, and extracts were immunoblotted with anti-caspase-9 (Casp-9^Pro^ and Casp-9^Cleaved^), anti-caspase-3 (Casp-3^Pro^ and Casp-3^Cleaved^), and anti-tubulin antibodies. **(G-H)** For quantification, the caspase cleavage ratio was calculated by dividing the cleaved band intensity by the pro-caspase band intensity. Each point represents the mean ratio ±SD from 3 experiments. AU = Arbitrary Units. *p<0.05; **p<0.01; ***p<0.001 (ANOVA, Tukey post-hoc tests).

We next sought to measure the effects of JMY and WHAMM further upstream in the caspase cascade by examining the expression and cleavage of representative initiator and executioner caspase proteins via immunoblotting. Control DMSO-treated parental, JMY^KO^, and WHAMM^KO^ cells expressed equivalent levels of inactive initiator procaspase-9 and inactive executioner procaspase-3 (Fig 4E and 4F), suggesting that the mutant cell lines did not have any inherent deficits in the steady-state abundance of these core components of the apoptotic machinery. However, upon etoposide treatment, the parental, JMY^KO^, and WHAMM^KO^ cell lines displayed very different kinetics in the conversion of inactive procaspases to cleaved caspases. Whereas HAP1 and eHAP cells began to show processing of procaspases to cleaved caspases by 3h of etoposide treatment, the cleaved caspases were generally not detectable in the JMY^KO^ and WHAMM^KO^ cells until 6h (Fig 4E and 4F). Even at the later timepoint, caspase-9 cleavage was 2-3-fold lower in JMY^KO^ cells and WHAMM^KO^ cells compared to their respective parental cell lines (Fig 4G and 4H). For caspase-3, cleavage was 10-fold less efficient in JMY^KO^ cells and 3-fold less efficient in WHAMM^KO^ cells relative to their parental cells (Fig 4G and 4H). The caspase-3 results were further validated by immunofluorescence microscopy, where 35% of parental cells stained positive for active caspase-3, compared to only 8% of the JMY^KO^ and 15% of the WHAMM^KO^ cells (S5 Fig). Together, these results show that following severe genotoxic stress, JMY and WHAMM are required for the rapid activation of caspase cleavage cascades.

Intrinsic pathways of apoptosis are characterized by the export of apoptogenic proteins from mitochondria into the cytosol upstream of the initiation of the caspase cascade. So we next examined if the loss of JMY or WHAMM affected the release of cytochrome *c* (cyto *c*), a protein that is maintained in the intermembrane space under normal conditions, but is exported to promote cytosolic assembly of the apoptosome and activation of caspase-9 during intrinsic apoptosis [40]. Consistent with the previously-observed amounts of AnnV-positive cells and levels of caspase cleavage/activation, >30% of parental cells exhibited diffuse cytosolic cyto *c* localization following 6h of etoposide treatment (Fig 5A-D). In contrast, the JMY^KO^ and WHAMM^KO^ cell lines had significantly lower proportions of cells with diffuse cyto *c* staining (Fig 5A-D). These findings collectively indicate that JMY and WHAMM are necessary for efficient intrinsic apoptotic responses during and/or prior to the accumulation of cyto *c* in the cytosol.

**Fig 5.**
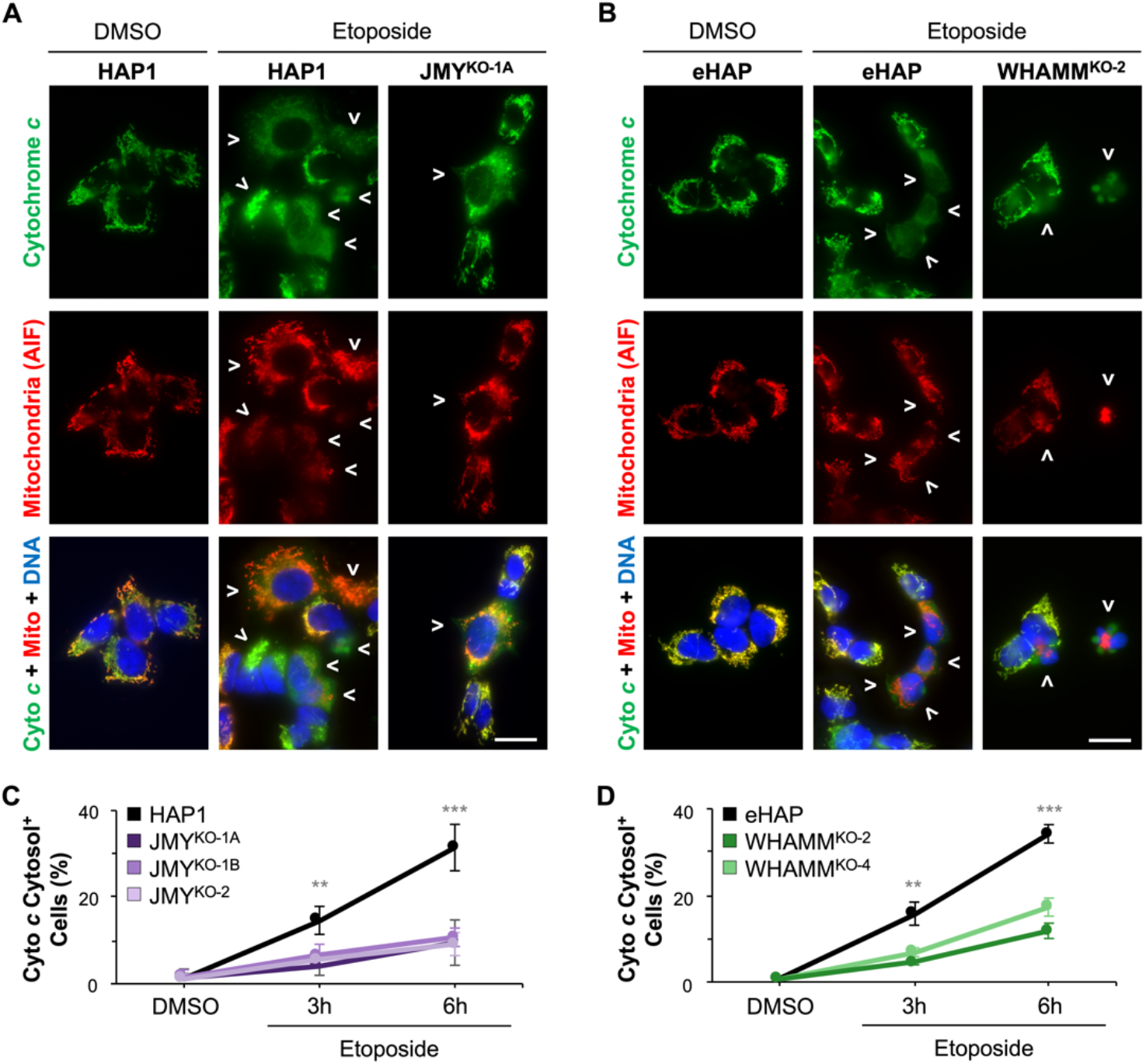
Cytochrome *c* release is delayed when JMY or WHAMM are not present. **(A-B)** HAP1, JMY^KO^, eHAP, and WHAMM^KO^ cells were treated with DMSO for 6h or 5 μM etoposide for 3h or 6h before being fixed and stained with a cytochrome *c* antibody (Cyto *c*; green), AIF antibody (Mito; red), and DAPI (DNA; blue). Images are from the 6h timepoint. Arrows highlight examples of diffuse cytosolic cyto *c* staining. Scale bar, 25μm. **(C-D)** The % of cells with cytosolic cyto *c* staining was calculated in ImageJ by counting cells that exhibited diffuse cytosolic rather than focused mitochondrial cyto *c* staining and dividing by the total number of DAPI-stained nuclei. Each point represents the mean % ±SD from 3-4 experiments (n = 533-757 cells per point in C; n = 629-856 cells per point in D). Significance stars refer to comparisons of parental to KO samples at the depicted timepoints. **p<0.01; ***p<0.001 (ANOVA, Tukey post-hoc tests).

### JMY transitions cells from p21-associated cell cycle arrest to p53-dependent cell death

To begin to understand the underlying mechanisms that give rise to the apoptotic pathway defects in JMY- and WHAMM-deficient cells, we sought to determine the importance of p53 in the death of parental HAP1 cells. First, to verify that HAP1 apoptosis is mediated by p53, we treated these parental cells with three independent siRNAs targeting p53 and measured the percentage of cells with AnnV staining. Compared to samples that were treated with a control siRNA, samples in which p53 levels were diminished contained significantly fewer AnnV-positive cells (Fig 6A and 6B). Moreover, greater degrees of p53 knockdown resulted in larger reductions in apoptosis, as the amount of p53 protein was positively correlated with the percentage of AnnV-positive cells (Fig 6C). The latter trend closely paralleled the phenotype of JMY-depleted cells (Fig 3D). These results demonstrate that the majority of etoposide-induced apoptosis in HAP1 cells requires p53.

**Fig 6.**
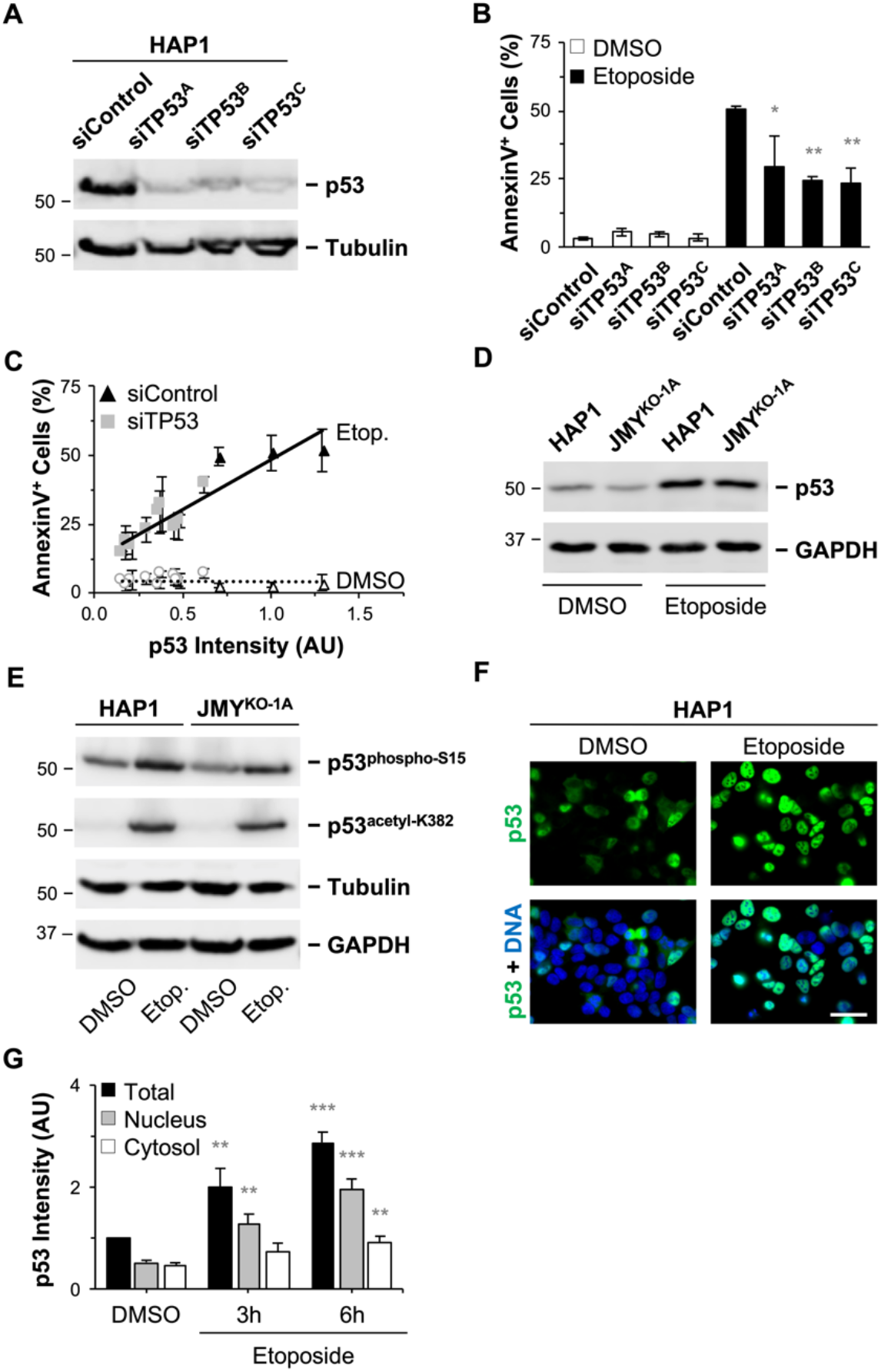
JMY is not required for p53 stabilization, nuclear accumulation, phosphorylation, or acetylation. **(A)** HAP1 cells were treated with control siRNAs or independent siRNAs for the TP53 gene before immunoblotting with anti-p53 and anti-tubulin antibodies. **(B)** The cells were treated with DMSO or 5μM etoposide for 6h and stained with Alexa488-AnnV, PI, and Hoescht. The % of AnnV-positive cells was calculated and each bar represents the mean ±SD from 3 experiments (n = 2,403-5,237 cells per bar). **(C)** p53 band intensities on immunoblots were normalized to tubulin bands and plotted against the % of AnnV-positive cells. Each point represents the mean ±SD from 3 fields-of-view in a given experiment (n = 350-2,217 cells per point). The slope in the linear trendline regression equation for etoposide-treated samples (Y = 36.61X + 11.81) was significantly non-zero (p<0.001, R^2^=0.75). **(D-E)** HAP1 and JMY^KO^ cells were treated with DMSO or etoposide for 6h before immunoblotting with anti-p53, anti-p53^phospho-S15^, anti-p53^acetyl-K382^, anti-tubulin, and anti-GAPDH antibodies. **(F)** HAP1 cells were treated with DMSO or etoposide for 6h before being fixed and stained with a p53 antibody (green) and DAPI (DNA; blue). Scale bar, 50μm. **(G)** Nuclear and cytosolic anti-p53 fluorescence intensities were measured in ImageJ and the total combined p53 intensity was normalized to the DMSO sample. Each bar represents the mean ±SD from 2 experiments (n = 149-151 cells per timepoint). Significance stars refer to comparisons to the DMSO sample. AU = Arbitrary Units. *p<0.05, **p<0.01, ***p<0.001 (ANOVA, Tukey post-hoc tests).

The observation that HAP1 cells undergo apoptosis in a p53-dependent manner, combined with the finding that JMY can enhance the expression of the pro-apoptotic p53 target *BAX* [24, 29], led us to next investigate the specific influence of JMY on p53 functions. Since apoptosis is typically accompanied by p53 protein stabilization, its phosphorylation on serine-15, acetylation on lysine-382, and re-localization into the nucleus [25, 26], we examined these properties. At steady-state, HAP1 and JMY^KO^ cells expressed similar amounts of p53 protein (Fig 6D), indicating that the loss of JMY did not cause an overall decrease in p53 abundance. Treatment of HAP1 cells with etoposide for 6h resulted in a clear increase in p53 protein levels (Fig 6D), its phosphorylation on serine-15 and acetylation on lysine-382 (Fig 6E), and its accumulation in the nucleus in the majority of cells (Fig 6F and 6G). Similar phenotypes were also observed in JMY^KO^ cells (Fig 6D and 6E), suggesting that JMY functions downstream of such pro-apoptotic changes in p53 stability and modification.

In response to genotoxic damage, nuclear p53 alters transcription and can trigger cell cycle arrest, DNA repair, apoptosis, senescence, and other stress responses [41, 42]. Because a proliferation arrest is usually an early response to DNA damage, we compared the growth and death rates of HAP1 and JMY^KO^ cultures after exposure to etoposide. For these experiments, we treated each culture with DMSO or etoposide for 6h, removed the solvent or drug and replaced them with fresh media, and then quantified the numbers of total cells (live and dead) at regular intervals up to a 48h endpoint. While HAP1 and JMY^KO^ cells multiplied at equivalent rates after treatment with DMSO, both cell types stopped proliferating after treatment with etoposide (Fig 7A). In contrast, when apoptotic cell quantities were measured using AnnV staining, the proportion of cells undergoing apoptosis was significantly higher in the HAP1 samples at every timepoint (Fig 7B). For parental samples, >35% were AnnV-positive by 6h, 50% were apoptotic by approximately 10h, and the frequency of apoptosis had reached about 90% of cells by 48h (Fig 7B). JMY^KO^ samples never hit the 50% apoptosis mark, as only 35% were AnnV-positive at 48h (Fig 7B). Thus, without JMY, cells remain stuck in an arrested state and fail to shift into a proper death signaling mode.

**Fig 7.**
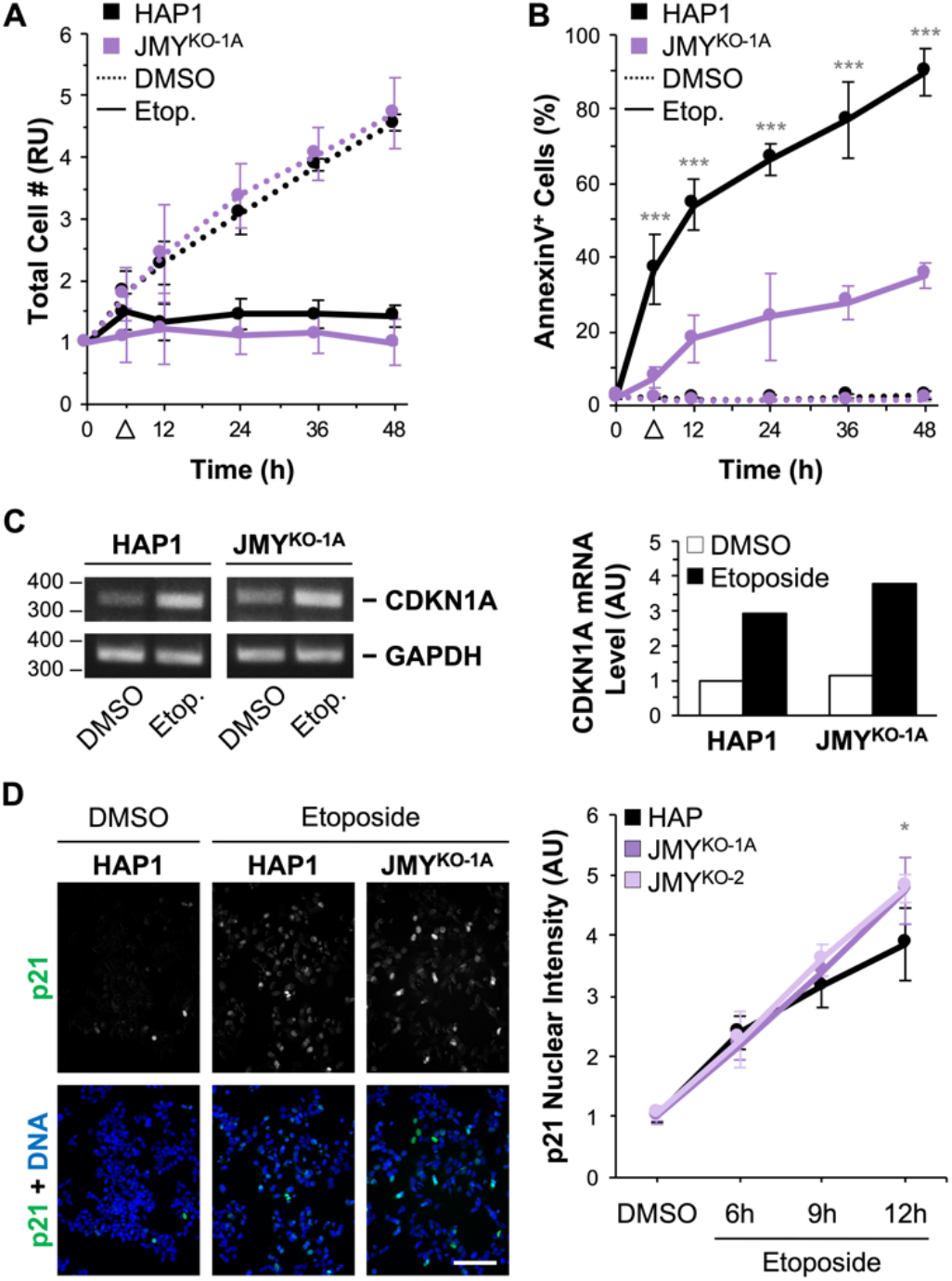
JMY-knockout cells undergo a prolonged p21-associated cell cycle arrest instead of rapid death following DNA damage. **(A-B)** HAP1 and JMY^KO^ cells were treated with DMSO or 5μM etoposide for 6h before washout (arrowhead) and replacement with regular media. Samples were stained with Alexa488-AnnV, PI, and Hoescht at the indicated time points. In (A), the total # of cells (live and dead) was counted in ImageJ and normalized to the # at 0h for each sample. In (B), the % of AnnV-positive cells was calculated and each point represents the mean ±SD from 3 experiments (n = 487-3,036 cells per point for DMSO; n = 472-995 cells per point for Etoposide). Significance stars refer to comparisons among the etoposide-treated samples. **(C)** HAP1 and JMY^KO^ cells were treated with DMSO or etoposide for 6h before collecting RNA and performing RT-PCR with primers for *CDKN1A* and *GAPDH*. Agarose gel band intensities were quantified in ImageJ, and values for *CDKN1A* were normalized to *GAPDH* and plotted in the adjacent bar graph. AU = Arbitrary Units. **(D)** Cells were treated with DMSO for 12h or etoposide for 6h, 9h, or 12h before being fixed and stained with a p21 antibody (green) and DAPI (DNA; blue). Images are from the 9h timepoint. Nuclear p21 fluorescence intensity was measured in ImageJ, and each bar represents the mean ±SD from 3 fields-of-view in a representative experiment (n = 504-1,946 cells per bar). Scale bar, 100μm. *p<0.05; ***p<0.001 (ANOVA, Tukey post-hoc tests).

One of the key p53 targets that triggers cell cycle arrest is *CDKN1A*, which encodes the cyclin-dependent kinase inhibitor, p21 [43]. To determine if *CDKN1A* expression is affected by *JMY* inactivation in the absence or presence of etoposide, we used RT-PCR for comparing *CDKN1A* transcript levels. Consistent with the proliferation assays described above, *CDKN1A* was expressed at low levels in DMSO-treated HAP1 and JMY^KO^ cells, and significantly upregulated in both cell lines following a 6h exposure to etoposide (Fig 7C). Similarly, small amounts of p21 protein were found in HAP1 and JMY^KO^ cells at steady-state (Fig 7D), while treatment with etoposide resulted in clear increases in the abundance of nuclear p21 in both sets of cells (Fig 7D). Collectively, these experiments suggest that JMY specifically drives a cell suicide program and is not required for several other aspects of nuclear p53 function, including the transcriptional responses that lead to cell cycle arrest.

### Expression of the small G-protein RhoD is turned on in JMY knockout cells

While inactivation of *JMY* did not prevent *CDKN1A* upregulation or cell cycle arrest, it could still impact other aspects of transcriptional programming. Indeed, one plausible explanation for the apoptotic defects in JMY^KO^ cells could be that they possess different gene expression patterns than normal cells such that they are less ‘equipped’ to die. To characterize the transcriptomic changes that took place upon knocking out *JMY*, we performed differential gene expression analyses using RNA-sequencing (RNA-seq) on HAP1 cells and on one of the JMY knockout cell lines (JMY^KO-1A^). When comparing the JMY^KO-1A^ cells to parental HAP1 cells, <0.36% of protein-coding genes displayed expression differences of at least 2-fold and with a significance q-value of <0.05 (Fig 8A and 8B). Expression of the genes encoding p53, caspases, Bcl-2 family members, other key modulators of apoptosis, or JMY-interacting proteins such as p300, Strap, and Mdm2 were not significantly different (S2 Table). In addition, genes for WASP-family members and other actin nucleation factors were not substantially changed (S2 Table). Therefore, mutating *JMY* does not appear to adversely affect the expression of canonical apoptosis regulators or factors that are structurally or functionally related to JMY.

**Fig 8.**
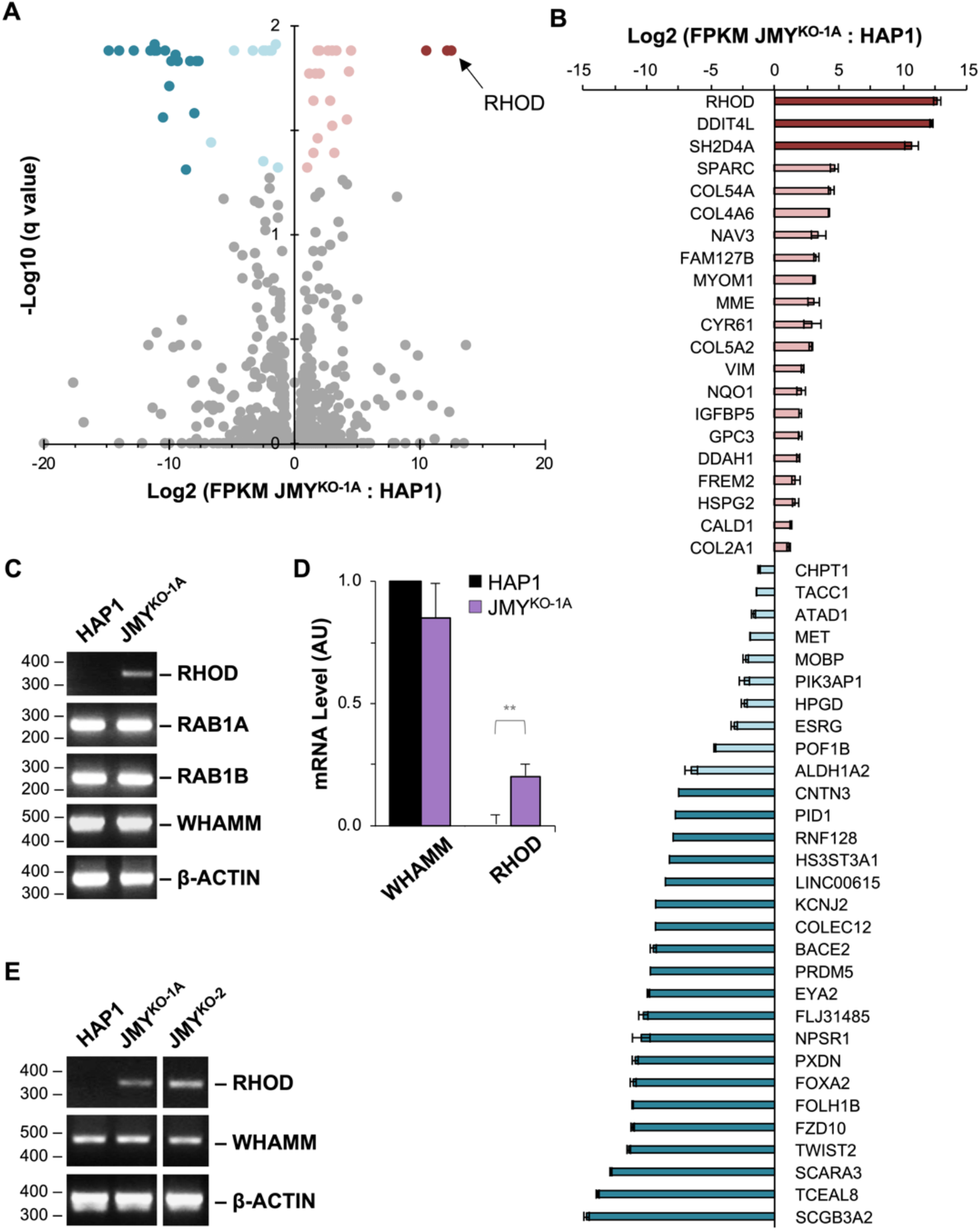
Expression of RhoD is turned on in JMY-knockout cells. **(A)** RNA collected from HAP1 and JMY^KO^ cells was subjected to mRNA sequencing analysis. A volcano plot representing FPKM (Fragments Per Kilobase of transcript per Million mapped reads) values of all individual genes for JMY^KO-1A^ vs HAP1 cells is plotted against the −log10(q value). Genes with expression differences of at least 2-fold and with a significance q-value of <0.05 are shown in dark red (turned on), pink (up-regulated), light blue (down-regulated), or dark blue (turned off), while genes that do not meet these criteria are depicted in gray (unchanged). Each point represents the mean value from 3 independent RNA samples per genotype. **(B)** FPKM values of individual genes with significant expression differences in (A) are shown. Each bar represents the mean ratio ±SD from 3 experiments. **(C)** RNA from HAP1 and JMY^KO^ cells was subjected to RT-PCR using primers to *RHOD*, *RAB1A*, *RAB1B*, *WHAMM*, and *β-ACTIN*, and visualized on an agarose gel. **(D)** Agarose gel band intensities were quantified in ImageJ, and values for *RHOD* and *WHAMM* were normalized to *β-ACTIN*. AU = Arbitrary Units. Each bar represents the mean ±SD from 3 experiments. **(E)** cDNAs from HAP1, JMY^KO-1A^, and JMY^KO-2^ cells were subjected to RT-PCR and visualized on an agarose gel. **p<0.01 (t-test).

Interestingly, however, of the 51 genes exhibiting significant differences in expression, the most upregulated gene in the JMY^KO^ line was *RHOD* (Fig 8A and 8B), which encodes a small G-protein that was previously shown to interact with WHAMM [44, 45]. Transcript levels for other small G-proteins were virtually the same between parental and KO cells (S2 Table), implying that the change for *RHOD* might be functionally meaningful. Moreover, RT-PCR experiments confirmed that *RHOD* transcript levels were almost undetectable in HAP1 cells and significantly turned on in the JMY^KO^ cells (Fig 8C and 8D). The expression of *WHAMM* and *RAB1A/RAB1B*, the latter of which encode small G-proteins known to bind WHAMM directly [46], were unchanged between the parental and JMY^KO^ cells (Fig 8C; S2 Table). RT-PCR further indicated that the independently derived *JMY* knockout cell line harboring a mutation in a different exon (JMY^KO-2^) also showed increased *RHOD* transcript levels (Fig 8E). These experiments reveal that although inactivating *JMY* may not cause any obvious compensatory changes in the expression of actin nucleation factors, it can result in the upregulation of a G-protein known to interact with JMY’s closest homolog, WHAMM.

### RhoD promotes cell survival in the absence or presence of JMY

RhoD is a Rho-family GTPase that regulates many cellular functions, including actin assembly during filopodia formation, cell migration, endosome dynamics, and Golgi trafficking [44–55]. Additionally, RhoD appears to participate in other processes that affect cell proliferation, such as cell cycle regulation and cytokinesis [48, 56]. Since JMY is involved in proteostasis and genotoxic stress responses through its activities in autophagy and apoptosis, we reasoned that the increase in *RHOD* expression in JMY^KO^ cells might be part of a compensatory pro-survival mechanism that allows the cells to better tolerate the loss-of-function mutation in *JMY*. To explore this possibility, we targeted RhoD for depletion using two independent siRNAs in JMY^KO^ cells. RT-PCRs verified that each RhoD siRNA reduced *RHOD* transcript levels (Fig 9A). DMSO-or etoposide-treated JMY^KO^ cells were then subjected to AnnV, PI, and Hoescht staining for measuring apoptosis and for evaluating necrosis (Fig 9B). Compared to control siRNA-treated samples, *RHOD*-depleted samples that were exposed to just DMSO showed modest increases in the fraction of cells with AnnV staining (Fig 9B and 9C), consistent with the possibility that cells are slightly more prone to apoptosis when *RHOD* levels are decreased. For the samples that were treated with etoposide, the low frequency of AnnV staining of JMY^KO^ cells was equivalent whether or not *RHOD* was silenced (Fig 9B-D). Since diminishing *RHOD* expression in these experiments did not restore apoptosis to normal levels, the intrinsic apoptotic defects in JMY^KO^ cells must not be due simply to the upregulation of *RHOD*. Perhaps more importantly, the percentage of AnnV-negative cells that were PI-positive, or considered to be necrotic, was significantly higher when *RHOD* was depleted (Fig 9B and 9C). This latter result supports the idea that RhoD normally promotes cell survival.

**Fig 9.**
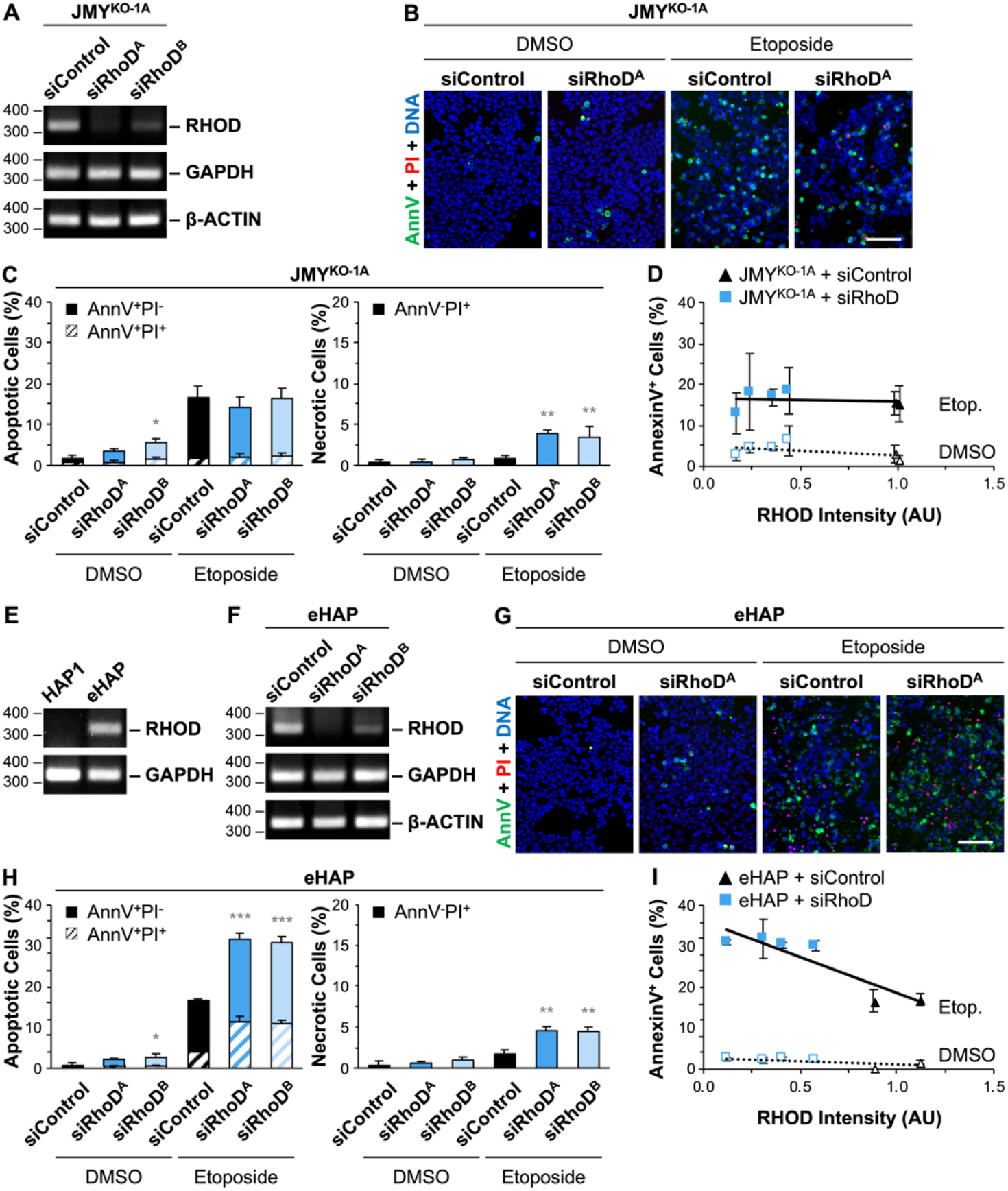
Transient depletion of RhoD results in greater levels of apoptotic and necrotic cell death. **(A)** JMY^KO^ cells were treated with control siRNAs or independent siRNAs for the RHOD gene before performing RT-PCRs with primers to *RHOD*, *GAPDH*, and *β-ACTIN*. **(B)** Cells were treated with DMSO or 5μM etoposide for 6h and stained with Alexa488-AnnV (green), PI (red), and Hoescht (blue). Scale bar, 100μm. **(C)** The % of AnnV-positive cells was calculated and displayed as the fraction of AnnV-positive/PI-negative (AnnV^+^PI^−^) or AnnV/PI double-positive (AnnV^+^PI^+^) cells. Significance stars refer to comparisons of total AnnV^+^ counts for siControl vs siRhoD samples. The % of AnnV-negative/PI-positive (AnnV^−^PI^+^) necrotic cells was also quantified. Significance stars refer to comparisons of siControl vs siRhoD samples. Each bar represents the mean ±SD from 3 experiments (n = 3,166-4,998 cells per sample). **(D)** *RHOD* band intensities were normalized to *β-ACTIN* and plotted versus the % of AnnV-positive cells. Each point represents the mean ±SD from 3 fields-of-view in a given experiment (n = 670-1,919 cells per point). **(E-F)** HAP1 and eHAP cells were subjected to RT-PCRs as in (A). **(G)** The cells were treated and stained as in (B). **(H)** Apoptosis and necrosis quantifications were performed as in (C). Each bar represents the mean ±SD from 3 experiments (n = 3,622-4,948 cells per sample). **(I)** *RHOD* levels were plotted against the % of AnnV-positive cells as in (D) (n = 908-1,728 cells per point). The slopes in the linear trendline regression equations for DMSO-treated samples (Y = −3.46X + 6.47) (p<0.01, R^2^=0.37) and etoposide-treated samples (Y = −43.48X + 90.28) (p<0.001, R^2^=0.75) were significantly non-zero. *p<0.05, **p<0.01, ***p<0.001 (ANOVA, Tukey post-hoc tests).

Because of the complexities in studying RhoD activities during apoptosis in JMY^KO^ cells, which appear to be fundamentally defective in their intrinsic apoptotic responses, we next wanted to characterize the effects of transiently depleting or permanently inactivating *RHOD* in otherwise healthy JMY-proficient cells. While *RHOD* transcript is below the limit of detection in the HAP1 cell line, it is expressed at measurable levels in eHAP cells (Fig 9E). So eHAP samples were treated with siRNAs for depleting *RHOD* (Fig 9F), exposed to DMSO or etoposide, and stained with AnnV and PI to assess the proportion of apoptotic and necrotic cells (Fig 9G). For the DMSO-treated populations, apoptotic death rose from 1% in cells receiving the control siRNA to 5% in cells receiving either of the RhoD siRNAs (Fig 9G and 9H), suggesting that RhoD depletion in healthy cells leads to a higher incidence of death under normal growth conditions. For the etoposide-treated samples, *RHOD* silencing caused apoptosis to occur in approximately 80% of eHAP cells, frequencies which were approximately double what was observed in cells treated with a negative control siRNA (Fig 9G and 9H). Moreover, the amount of RhoD transcript inversely correlated with the percentage of AnnV-positive cells (Fig 9I). Necrotic death frequencies also rose, in this case from 2% in the presence of *RHOD* to 5% when it was silenced (Fig 9H). Thus, decreasing *RHOD* expression can increase stress-induced apoptosis and necrosis when normal JMY and p53 responses are present, further strengthening the conclusion that RhoD can play a basic pro-survival role in cells.

To assess the impact of a permanent loss of *RHOD* on cell survival and death, we next derived two independent RhoD^KO^ cell lines from parental eHAP cells (Fig 10A and S2 Fig). RT-PCRs verified that wild type RhoD mRNA was absent in the mutant cells (Fig 10A). Similar to other knockout cells, each of the RhoD^KO^ cell lines had low levels of DNA damage when exposed to DMSO, but more numerous DNA breaks when incubated with etoposide, as evidenced by increased levels of γH2AX and its accumulation in nuclear foci (S3 Fig). Consistent with the transient depletion studies in etoposide-treated samples, early apoptotic, late apoptotic, and necrotic death were all more common in RhoD^KO^ cells than in eHAP cells (Fig 10B). Additionally, in dose-response experiments, the RhoD^KO^ cell lines showed etoposide-dependent increases in cell death that were greater than those of parental cells, with approximately twice as many apoptotic RhoD^KO^ cells observed at each etoposide concentration (Fig 10C). These experiments demonstrate that the permanent loss of RhoD makes cells more prone to dying during normal culture conditions and also enhances their apoptotic responses following acute DNA damage.

**Fig 10.**
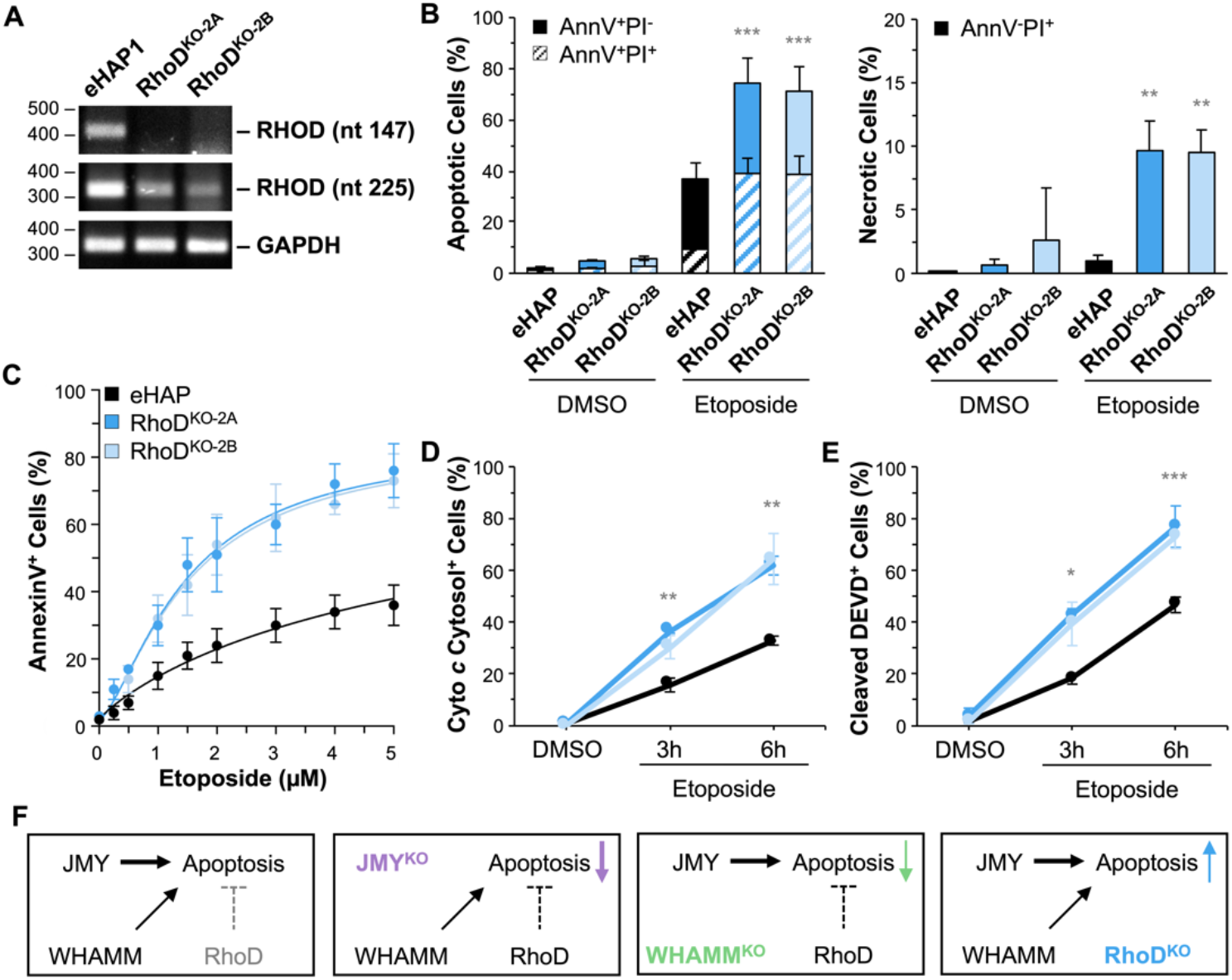
RhoD-knockout cells display enhanced apoptotic responses following DNA damage. **(A)** RNA from eHAP and RhoD^KO^ cells was subjected to RT-PCR using primers to *RHOD* (forward primer: nt 124-147; reverse primer: nt 523-545), *RHOD* (forward primer: nt 225-247; reverse primer nt 523-545), and *GAPDH*. Nucleotide 147 is predicted to be absent in mRNA from both knockouts, whereas nucleotide 247 should be present in both, albeit reflecting transcript levels that are lower than the parental control due to nonsense-mediated decay. Bands from this gel are also shown in Fig 9E. **(B)** eHAP and RhoD^KO^ cells were treated with DMSO or 5μM etoposide for 6h and stained with Alexa488-AnnV, PI, and Hoescht. The % of apoptotic and necrotic cells were quantified and each bar represents the mean ±SD from 3 experiments (n = 4,713-9,322 cells per sample). **(C)** eHAP and RhoD^KO^ cells were treated with a range of etoposide concentrations for 6h and then stained with Alexa488-AnnV and Hoescht. The % of AnnV-positive cells was calculated and each point represents the mean ±SD from 3-6 fields-of-view pooled from 1-2 experiments (n = 2,309-4,356 cells per experiment). Nonlinear regressions were performed with a baseline set to 0.02 and a maximum response set to 0.85. The EC50 is significantly different for parental vs KO samples (EC50: eHAP = 6.8 μM; RhoD^KO^ = 1.4-1.5 μM, p<0.001). **(D)** Cells were treated with DMSO or 5 μM etoposide for 3 or 6h, fixed, and stained with probes to detect cyto *c*, AIF, and DNA. The % of cells with cytosolic cyto *c* staining was calculated and each point represents the mean ±SD from 3 experiments (n = 480-843 cells per point). Significance stars refer to comparisons of parental to KO samples at the depicted timepoints. **(E)** Cells were stained with caspase-3/7 green detection reagent, and the % of cleaved DEVD-positive cells was calculated by counting cells that exhibited green nuclear fluorescence and dividing by the total number of Hoescht-stained nuclei. Each point represents the mean ±SD from 3 experiments (n = 2,329-5,804 cells per point). *p<0.05, **p<0.01, ***p<0.001 (ANOVA, Tukey post-hoc tests). Representative images and immunoblots appear in (S6 Fig). **(F)** Model for JMY, WHAMM, and RhoD in apoptosis. Line thickness reflects different degrees of apoptotic phenotypes, dashed lines represent conditional roles for RhoD, gray text indicates a lack of RhoD expression.

Since JMY and WHAMM are necessary for an efficient pathway of cyto *c* release, initiator caspase cleavage, and executioner caspase activation, we tested if *RHOD* inactivation affected these aspects of intrinsic apoptosis. Following 3h and 6h of etoposide treatment, compared to the eHAP parental cell line, the RhoD^KO^ samples had significantly higher percentages of cells with diffuse cyto *c* localization (Fig 10D and S6 Fig), greater levels of initiator caspase-9 cleavage (S6 Fig), more cells staining positive for cleaved caspase-3 (S6 Fig), and more cells undergoing cleavage of DEVD-containing substrates (Fig 10E and S6 Fig). Together, the above results show that knocking out *RHOD* has the opposite apoptotic phenotypes of knocking out *JMY* or *WHAMM*. Thus, while JMY and WHAMM are WASP-family proteins that have important pro-apoptotic roles in intrinsic pathways of cell death, such functions may be constrained by pro-survival or anti-apoptotic activities of the small G-protein RhoD.

## Discussion

The polymerization, organization, and turnover of actin filaments have been thoroughly characterized during many cellular functions that maintain viability [9, 10]. However, the contributions of cytoskeletal factors to apoptotic cell death are not well understood. Perhaps the best known cytoskeletal features of apoptosis are the actin filament rearrangements and disassembly that accompany changes in cell adherence and morphology [7, 8]. Actin itself can be cleaved by caspases [57, 58], resulting in non-polymerizable fragments that localize to mitochondria and promote apoptotic phenotypes when expressed in cells [59–61]. Importantly, the actin turnover machinery has also been implicated in controlling apoptosis at multiple stages. Cofilin, which depolymerizes actin filaments [62], influences the early stages of staurosporine-induced apoptosis, as it translocates to mitochondria and increases the release of cytochrome *c* [63–65]. Additionally, active cofilin interacts with p53 to affect its localization to the nucleus and mitochondria to promote apoptosis [66]. Actin is also recruited to mitochondria at around the time of mitochondrial permeabilization [67, 68]. During the later stages of apoptosis, the F-actin-severing protein gelsolin [69] is cleaved by caspases, resulting in an N-terminal fragment with unregulated severing activity that drives depolymerization [70]. This gelsolin fragment additionally releases DNase I from its interaction with monomeric actin to allow DNase I-mediated chromatin fragmentation [71]. In contrast to these described roles for actin disassembly proteins in apoptosis, less is understood about the actions of actin assembly proteins during cell death. Our results reveal intrinsic apoptotic functions for JMY, WHAMM, and RhoD, and create a framework for understanding how actin nucleation factors and small G-proteins control pro-apoptotic and pro-survival processes.

WASP-family proteins have well-recognized activities in many vital cellular functions, but the extent to which they participate in cell death pathways is relatively uncharacterized. WAVE1 has been one of the most studied members, as it can influence the localization or modification of Bcl-2-family proteins. Transient overexpression and depletion experiments suggest that WAVE1 limits mitochondrial permeabilization in a cell type dependent manner [20–23]. But in our study, permanently knocking out the gene encoding WAVE1 in BCR-ABL-immortalized human cells increased the frequency of apoptosis in only 1 of 3 experiments. So the degree to which WAVE1 acts as an anti-apoptotic factor requires further investigation.

In addition to WAVE1, JMY has a reported role in apoptosis. JMY was discovered as a cofactor that enhances the transcriptional activity of p53 [24], and subsequent RNAi studies indicated that JMY has pro-survival [32] or pro-death [27, 29, 72] functions under different experimental circumstances. Our work advances the understanding of its pro-death role by demonstrating that cells with permanent deletions or transient depletions of JMY exhibit significant defects in apoptosis following DNA damage. In fact, inactivation of each WASP-family gene revealed that only cells lacking JMY, or its closest homolog WHAMM, had substantial deficiencies in their intrinsic apoptotic responses. WHAMM did not have any previously reported roles in cell death, so the observation that WHAMM is important for efficient apoptosis adds a third WASP-family member to the repertoire of actin assembly factors that function in programmed cell death.

WHAMM and JMY comprise a subgroup within the WASP family, are approximately 50% similar to one another [37], and both promote anterograde membrane transport [37, 46, 73] and autophagy [32, 74, 75]. Although the two homologous proteins participate in similar cellular processes, they exhibit some key differences. WHAMM has a conventional C-terminal WWCA domain that activates the Arp2/3 complex [37], whereas JMY has an extended WWLWCA domain that includes a Linker which enables it to directly nucleate linear actin filaments in addition to assembling branched networks via the Arp2/3 complex [76]. WHAMM localizes to the ER and *cis*-Golgi region and binds microtubules to mediate membrane tubulation and transport [37, 46, 77], while JMY acts later in the secretory pathway to promote trafficking away from the *trans*-Golgi [73]. WHAMM and JMY also both function in multiple aspects of autophagy. WHAMM binds to the phospholipid PI(3)P and activates the Arp2/3 complex to somehow accelerate lipidation of LC3-family proteins during autophagosome biogenesis [75]. WHAMM additionally triggers autophagosome movement via actin ‘comet’ tails [74] and can further drive autolysosome remodeling [78]. JMY binds to LC3 and polymerizes actin at autophagosomes during their maturation [32] and can also assemble actin comet tails [28]. While it is reasonable to assume that the functions of WHAMM and JMY in secretion and autophagy help sustain cells during normal growth and allow them to adapt to conditions of nutritional stress, the fact that both factors also enable cell death indicates that they function as pivotal players in the cellular responses to multiple other stressors including genotoxic damage.

Following etoposide-induced DNA breaks, parental HAP1 cells executed an immediate p53-dependent cell death program. In contrast, JMY or WHAMM knockout cells showed significant reductions in the terminal apoptosis phenotypes of phosphatidylserine externalization, membrane permeabilization, and nuclear fragmentation. Importantly, both sets of knockout cells also displayed deficiencies and/or kinetic delays in earlier apoptotic processes, including the mitochondrial export of apoptogenic proteins, caspase activation, and effector caspase cleavage of target proteins. These results indicate that JMY and WHAMM are key contributors to a rapid, intrinsic, p53-mediated cell death pathway. They further suggest that JMY and WHAMM each function after the accumulation of DNA damage but before or during the step of mitochondrial permeabilization. It is also important to note that one or both of these factors may additionally accelerate later processes such as apoptosome assembly and caspase cleavage/activation.

Under normal cell culture conditions, JMY is found predominantly in the cytosol [27, 28], but in response to DNA damage, it accumulates in the nucleus while still maintaining some cytosolic presence [29, 30]. At steady-state, WHAMM is mostly associated with cytoplasmic membrane-bound organelles, although it displays a nuclear localization in a small fraction of cells [37]. Such observations are consistent with the existence of both cytoplasmic and nuclear roles for these proteins in pro-apoptotic pathways. While the loss of JMY or WHAMM results in similar types of apoptotic deficiencies, the extent to which apoptosis would proceed in cells lacking both homologs remains to be determined. Nevertheless, the more severe phenotypes observed when *JMY* is mutated suggest that JMY is a more prominent player in cell death.

JMY was previously shown to interact with the stress-response protein Strap, the acetyltransferase p300, and p53 [24, 31]. p300 is one of many cofactors known to modify p53 to promote the transcription of genes necessary for initiating cell cycle arrest and stimulating pro-repair or pro-death pathways in response to cell damage [26, 79–84]. While the precise function of JMY in a complex with Strap, p300, and p53 is unknown, overexpression and depletion studies indicate that JMY enhances transcription of the pro-apoptotic gene *BAX*. JMY overexpression is not sufficient to increase *BAX* expression, but co-overexpression of JMY with p53, or with p53 plus p300, augments *BAX* transcription in several epithelial cell contexts [24, 29, 72]. Consistent with JMY being important for *BAX* upregulation, introduction of a JMY siRNA into p53-overexpressing cells limits the induction of *BAX* [29]. Our current work provides additional insight into how JMY impacts p53 functions during genotoxic stress.

p53 shuttles between the cytoplasm and nucleus, and although it has pro-apoptotic functions in the cytosol [85, 86], its most extensively characterized activities are as a nuclear transcription factor [42, 87, 88]. In our experiments, DNA damage-induced increases in the abundance and nuclear accumulation of p53 occurred in the presence or absence of JMY. Phosphorylation and acetylation of p53 was similar in both parental and knockout cells, indicating that JMY is not necessary for the typical pro-apoptotic changes in p53 stability, nuclear import, or posttranslational modification. Moreover, *JMY* inactivation had little effect on the transcriptional upregulation of *CDKN1A*, which encodes the key cell cycle inhibitor p21. In fact, JMY-deficient cells exhibited a prolonged cell cycle arrest in response to DNA damage, indicating that JMY is not necessary for the transcriptional changes that stop cell cycle progression. These results collectively support the idea that JMY directs a specific subset of p53-mediated transcriptional responses to genotoxic stress. We propose that one key function of nuclear JMY is to transition cells from expressing cell cycle arrest genes to transcribing pro-apoptotic genes.

Given the speed with which p53- and JMY-proficient parental cells undergo apoptosis, we hypothesized that they constitutively express a pool of potentially apoptotic factors that enable the cells to respond rapidly to DNA damage or other insults. To explore whether JMY knockout cells possessed altered gene expression patterns compared to their JMY-expressing parents, we used RNA-seq to measure steady-state transcript levels in both cell lines. While expression differences in canonical apoptogenic factors were not apparent, it was striking that the JMY knockout cells experienced a massive upregulation of *RHOD*, which encodes a small G-protein that was previously shown to interact with WHAMM [44, 45]. RhoD has wide-ranging functions in regulating actin dynamics at the plasma membrane, endosomes, and Golgi, and in influencing cell proliferation [44–55]. We reasoned that increased RhoD expression may be a compensatory pro-survival mechanism that allows cells to better tolerate a permanent loss of JMY. Consistent with this possibility, RhoD loss-of-function experiments result in more cell death – the opposite phenotype of JMY-or WHAMM-depleted cells. These findings reveal complex relationships among two actin nucleation factors and one small G-protein during cell death and survival (Fig 10F). The physical connections among RhoD, WHAMM, JMY, and actin during apoptosis, and the precise signaling activities of RhoD that modulate the cellular responses to genotoxic, nutritional, or other stresses require further investigation.

Continuing to define the roles of different components of the actin assembly and regulatory machinery in cellular adaptations to endogenous or exogenous stress could provide new avenues for understanding tumorigenesis and therapeutic interventions. All WASP-family members influence cell motility [13, 14], and elevated levels of nucleation factors are associated with increased metastasis [89, 90], indicating that such actin cytoskeletal proteins have proto-oncogenic features. Similarly, the functions of WHAMM and JMY in autophagy [32, 74, 75] could enable cancer cells to better survive in diverse physiological environments and in response to chemotherapy [91, 92]. However, our current results describing apoptotic requirements for JMY and WHAMM support the idea that these factors also possess key tumor-suppressive features. *TP53* is well known as the most commonly mutated gene in human cancers, and mutations that inactivate p53 or other p53-associated signaling components have been linked to poor prognosis [93–95]. Interestingly, JMY expression also appears to be lost in several B-cell lymphomas and invasive carcinomas [96]. Given the new positions of JMY and WHAMM as important components of p53-dependent apoptotic pathways, a greater understanding of how their actin nucleation, proto-oncogenic, and tumor-suppressor activities are coordinated will likely shed further light on how programmed cell death mechanisms are impacted by the cytoskeleton.

## Materials and Methods

### Ethics statement

Research with biological materials in the Campellone Lab was approved by the UConn Institutional Biosafety Committee (#759C). This study did not include research with human subjects or live animals. Human cell lines were acquired from Horizon Genomics or the UC Berkeley cell culture facility as described below.

### Cell culture

Human cell lines are listed in S1 Table. HAP1 cells are nearly-haploid fibroblast-like cells that contain an immortalizing BCR-ABL fusion and a single copy of all chromosomes except for a heterozygous 30Mb fragment of chromosome 15, which is integrated within the long arm of chromosome 19 [33, 97]. This diploid portion encompasses 330 genes, including *WHAMM*. CRISPR/Cas9-engineered eHAP cells were derived from the HAP1 line and are fully haploid [35]. In the current study, CRISPR/Cas9-mediated recombination using guide RNAs against target genes resulted in frameshift and/or splicing mutations (S1 Table) which were confirmed by DNA sequencing (Horizon Genomics). HAP1 cells were used for mutagenesis of *JMY*, *WASL* (encoding N-WASP), *WASF1* (WAVE1), *WASF2* (WAVE2), *WASF3* (WAVE3), *BRK1* (WAVE Complex), *CCDC53* (WASH Complex), and *CTTN* (Cortactin), while eHAP cells were used for mutagenesis of *WHAMM* and *RHOD*. The latter cell lines were made in an eHAP background due to the *WHAMM* diploidy in HAP1 cells, and because under normal culture conditions *RHOD* transcript was detectable in eHAP but not HAP1 cells (Horizon Genomics). One of the *WHAMM* mutant cell lines (KO-2) was studied previously [75]. HAP1 cell derivatives were cultured in Iscove’s Modified Dulbecco’s Medium (IMDM) supplemented with 10% fetal bovine serum (FBS) and penicillin-streptomycin. HeLa and U2OS cells (UC Berkeley cell culture facility) were cultured in Dulbecco’s Modified Eagle Medium (DMEM) supplemented with 10% FBS and antibiotic-antimycotic. All cell lines were grown at 37°C in 5% CO_2_. All assays were performed using cells that had been in culture for 2-10 passages.

### RT-PCR and RNA-sequencing

For RT-PCRs, approximately 10^6^ cells were seeded in 6-well plates, and for RNA-seq 2.5×10^6^ cells in were seeded in 6cm dishes. After 24h of growth, cells were rinsed with phosphate buffered saline (PBS). RNA was isolated using TRIzol reagent (Ambion), followed by a chloroform extraction, isopropanol precipitation, and 75% ethanol wash before resuspending RNA in water. For RT-PCRs, RNA was reverse transcribed into cDNA using Superscript III RT (Invitrogen) and subsequently amplified using Taq polymerase (New England Biolabs) and primers listed in S3 Table. Primers were designed to amplify ~230-460bp products from each cDNA template. After agarose gel electrophoresis, band intensities for *CDKN1A*, *WHAMM*, and *RHOD* were quantified in ImageJ [98] and normalized to *β-ACTIN* and/or *GAPDH* bands for each sample. For RNA-seq, Illumina cDNA library preparation was based on the Illumina TruSeq Stranded mRNA sample preparation guide. Total reads from RNA-seq were aligned with TopHat, and differential expression was calculated with the CuffLinks/CuffDiff program. Gene expression values were given as Fragments Per Kilobase of transcript per Million mapped reads (FPKM), differential expression was calculated as the Log2(FPKM ratio JMY^KO-1A^: HAP1), and data from 3 independent experiments were merged. Volcano plots were generated by plotting the −Log10(q value) against Log2(FPKM ratio JMY^KO-1A^: HAP1). The data summarized in this publication will be deposited in NCBI’s Gene Expression Omnibus.

### Chemical treatments and siRNA transfections

Cells were treated with different concentrations of etoposide (Sigma Aldrich) diluted from an initial stock of 10mM in DMSO. Equivalent volumes of DMSO were used as controls. Prior to siRNA transfections, cells were grown in 6-well plates for 24h. For RNA interference experiments, cells were transfected with 40nM Sigma MISSION siRNAs (S3 Table) using RNAiMAX (Invitrogen), incubated in growth media for 24h, reseeded into 6-well plates and 24-well glass-bottom plates (MatTek), and incubated for an additional 24h. Cells cultured in 6-well plates were collected and processed for immunoblotting or RT-PCR, and cells cultured in 24-well plates were used for live fluorescence microscopy assays.

### Immunoblotting and quantification

Detached cells were collected from 6-well plates and combined with adherent cells in PBS containing EDTA, centrifuged, washed with PBS, and centrifuged again to ensure collection of all live and dead material. Cell pellets were resuspended in lysis buffer (20mM HEPES pH 7.4, 100mM NaCl, 1% Triton X-100, 1mM Na_3_VO_4_, 1mM NaF, plus 1mM PMSF, and 10μg/ml each of aprotinin, leupeptin, pepstatin, and chymostatin), diluted in SDS-PAGE sample buffer, boiled, centrifuged, and subjected to SDS-PAGE before transfer to nitrocellulose membranes (GE Healthcare). Membranes were blocked in PBS + 5% milk (PBS-M) before being probed with primary antibodies diluted in PBS-M at concentrations listed in S4 Table overnight at 4°C plus an additional 2-3h at room temperature. Membranes were rinsed twice with PBS and washed thrice with PBS + 0.5% Tween-20 (PBS-T). Membranes were then probed with secondary antibodies conjugated to IRDye-800, IRDye-680, or horseradish peroxidase diluted in PBS-M at concentrations listed in S4 Table. Membranes were again rinsed with PBS and washed with PBS-T. Blots were imaged using a LI-COR Odyssey Fc imaging system. Band intensities were determined using the Analysis tool in LI-COR Image Studio software and quantities of proteins-of-interest were normalized to tubulin and/or GAPDH loading controls.

### Apoptosis and caspase activation assays

For live fluorescence-based assays, approximately 2.5×10^5^ cells were seeded into 24-well glass-bottom plates and allowed to grow for 24h prior to DMSO or etoposide treatments. For apoptosis assays, Alexa488-AnnexinV (Invitrogen), Propidium Iodide (Invitrogen), and Hoescht (Thermo Scientific) were added directly to the media at concentrations listed in S4 Table and incubated for 15min at 37°C in 5% CO_2_. For caspase activation assays, CellEvent Caspase-3/7 Detection Reagent (Invitrogen) and Hoescht were added to the media at concentrations listed in S4 Table and incubated for 30min at 37°C in 5% CO_2_. All live imaging was performed at 37°C as described below.

### Immunostaining assays

For fixed cell immunofluorescence microscopy, approximately 2.5×10^5^ cells were seeded onto 12mm glass coverslips in 24-well plates and allowed to grow for 24h. After DMSO or etoposide treatments, cells were washed with PBS, and fixed in 3.7% paraformaldehyde (PFA) in PBS for 30min, washed, permeabilized with 0.1% TritonX-100 in PBS, washed, and incubated in blocking buffer (1% FBS + 1% bovine serum albumin (BSA) + 0.02% NaN_3_ in PBS) for a minimum of 15min. Cells were probed with primary antibodies diluted in blocking buffer at concentrations listed in S4 Table for 45min. Cells were washed and treated with AlexaFluor-488 or −555 conjugated goat anti-rabbit or goat anti-mouse secondary antibodies, DAPI, and/or AlexaFluor-488 conjugated phalloidin at concentrations listed in S4 Table for 45min, followed by washes and mounting in Prolong Gold anti-fade reagent (Invitrogen).

### Fluorescence microscopy

All live and fixed images were captured using a Nikon Eclipse T*i* inverted microscope with Plan Apo 100X/1.45, Plan Apo 60X/1.40, or Plan Fluor 20x/0.5 numerical aperture objectives using an Andor Clara-E camera and a computer running NIS Elements software. Live cell imaging was performed in a 37°C chamber (Okolab) using the 20X objective. Fixed immunofluorescence imaging was performed using the 100X or 60X objectives. All cells were viewed in multiple focal planes, and even when Z-series were captured (at 0.2-0.4 μM steps), each of the images presented in the Figures represent a single slice. All images were processed and/or analyzed using ImageJ software [98].

### Image processing and quantification

The ImageJ Cell Counter plugin was used for analyses of live apoptosis assays, live caspase-3/7 activation assays, fixed cytochrome *c* assays, and fixed cleaved caspase-3 assays by manually counting the total number of cells in the Hoescht or DAPI channel and the number of cells that were positive for AnnV fluorescence, AnnV fluorescence plus nuclear PI fluorescence, nuclear PI fluorescence, nuclear DEVD reporter fluorescence, diffuse cytosolic cytochrome *c* staining, or cleaved caspase-3 staining. For γH2AX analyses, the Threshold, Watershed, and Analyze tools were used in the DAPI channel to separate individual nuclei, and the ROI Manager Measure tool was used in the γH2AX channel to measure mean γH2AX fluorescence intensity per nucleus. Additionally, the Find Maxima and ROI Manager Measure tools were used in the γH2AX channel to count the number of γH2AX foci per nucleus.

### Reproducibility and statistics

All conclusions were based on observations made from at least 3 separate experiments and quantifications were based on data from 3-5 representative experiments, except where noted in the Figure Legends. To capture a breadth of apoptotic phenotypes, multiple fields-of-view containing cells at different densities were imaged. The sample size used for statistical tests was the number of times an experiment was performed, except where noted in the Legends. Statistical analyses were performed using GraphPad Prism software as stated in the Legends. Statistics on data sets with 3 or more conditions were performed using ANOVAs followed by Tukey’s post-hoc test unless otherwise indicated. P-values for data sets comparing 2 conditions were determined using unpaired t-tests unless otherwise noted. P-values <0.05 were considered statistically significant.

## Acknowledgements

We thank Bo Reese at the UConn Center for Genome Innovation for guidance with RNA-seq and analysis, Alyssa Coulter for assistance with RNA-seq, Vanessa Vlaun for help with microscopy quantification, Frida Zink for help processing knockout cells, and L.T. Bear for support with experimental design. We also thank Katrina Velle, Aoife Heaslip, and Campellone Lab members for their comments on this manuscript.

## Supporting Information Captions

**S1 Table. Cell Lines**.

**S2 Table. Expression of genes encoding actin nucleation factors, cell cycle arrest proteins, canonical apoptosis regulators, and representative small G-proteins**.

**S3 Table. RNA and DNA oligonucleotides**.

**S4 Table. Immunofluorescence and immunoblotting reagents**.

**S1 Fig. Cells lacking the WASP-family members JMY or WHAMM undergo less apoptosis**. **(A)** Parental (HAP1, eHAP) and WASP-family knockout (JMY^KO^, WHAMM^KO^, WAVE1^KO^) cells were treated with DMSO or 5μM etoposide for 6h and stained with Alexa488-AnnexinV (AnnV; green), Propidium Iodide (PI; red), and Hoescht (DNA; blue). Scale bar, 100μm. **(B)** Representative images show AnnV-negative/PI-negative (AnnV^−^PI^−^) non-apoptotic, AnnV-positive/PI-negative (AnnV^+^PI^−^) early apoptotic, AnnV/PI double-positive (AnnV^+^PI^+^) late apoptotic, or AnnV-negative/PI-positive (AnnV^−^PI^+^) necrotic cells. Scale bar, 25μm. **(C)** Arrows highlight examples of hoescht-stained DNA condensation and nuclear fragmentation. Scale bar, 25μm.

**S2 Fig. JMY, WHAMM, and RhoD knockout cell lines contain loss-of-function mutations derived from frameshifts or altered splicing**. **(A)** HAP1 cells were treated with guide RNAs to the first or second exon of the *JMY* gene. A 17bp deletion in JMY^KO-1A^, a 10bp deletion in JMY^KO-1B^, and a 2bp deletion in JMY^KO-2^ resulted in premature stop codons. **(B)** eHAP cells were treated with guide RNAs to the second or fourth exon of the *WHAMM* gene. A 10bp deletion in WHAMM^KO-2^ and a 7bp deletion in WHAMM^KO-4^ resulted in premature stop codons. **(C)** eHAP cells were treated with guide RNAs to the second exon of the *RHOD* gene. A 22bp deletion in RhoD^KO-2A^ is predicted to result in defective splicing (not shown) and/or a frameshift (shown), while a 1bp deletion in RhoD^KO-2B^ results in a simple frameshift and premature stop codon.

**S3 Fig. Parental and knockout cell lines exhibit high levels of γH2AX expression and clustering in response to DNA damage**. **(A)** HAP1, eHAP, and knockout cell lines were treated with DMSO or 5μM etoposide for 6h before immunoblotting with anti-γH2AX and anti-tubulin antibodies. **(B)** HAP1 and JMY^KO^ cells were treated with DMSO or etoposide, fixed, and stained with a γH2AX antibody (red), Alexa488-phalloidin (F-actin; green), and DAPI (DNA; blue). Representative images show increased nuclear γH2AX foci upon etoposide treatment. Scale bar, 25μm. **(C)** Nuclear γH2AX fluorescence intensity was calculated using ImageJ (n = 48-70 nuclei per sample from a representative experiment). **(D)** The number of γH2AX foci per nucleus was determined using ImageJ (n = 54-81 nuclei per sample from a representative experiment). Significance stars are in reference to the etoposide treated eHAP cell line. Fewer γH2AX foci were observed for etoposide-treated RhoD^KO^ cells because of the loss of some dead cells prior to fixation to the slide. *p<0.05 (ANOVA, Tukey post-hoc tests).

**S4 Fig. Transient JMY depletion in multiple cell lines results in less apoptosis following DNA damage**. **(A)** U2OS or HeLa cells were treated with control siRNAs or independent siRNAs for the JMY gene before immunoblotting with anti-JMY and anti-tubulin antibodies. **(B)** The cells were treated with DMSO or 5μM etoposide for 6h and stained with Alexa488-AnnV (green), PI (red), and Hoescht (blue). Scale bar, 100μm. **(C)** The % of AnnV-positive cells was calculated and each bar represents the mean ±SD from 3 experiments (U2OS: n = 476-704 cells per bar; HeLa: 615-866 cells per bar). Significance stars refer to comparisons to the siControl samples. **(D)** JMY band intensities on immunoblots were normalized to tubulin bands and plotted against the % of AnnV-positive cells. Each point represents the mean ±SD from 3 images in a given experiment (U2OS: n = 69-342 cells per point; HeLa: n = 85-440 cells per point). The linear trendline regression equations for etoposide-treated samples (U2OS: Y = 38.31X + 3.86; HeLa: Y = 35.16X + 7.13) were significantly non-zero (p<0.001, R^2^>0.74). ***p<0.001 (ANOVA, Tukey post-hoc tests).

**S5 Fig. Caspase-3 cleavage is inefficient in JMY- and WHAMM-knockout cells**. **(A-B)** HAP1, JMY^KO^, eHAP, and WHAMM^KO^ cells were treated with DMSO or 5 μM etoposide for 6h before being fixed and stained with a cleaved caspase-3 antibody (Casp-3^Cleaved^; red), Alexa488-phalloidin (F-actin; green), and DAPI (DNA; blue). Scale bar, 25μm. The % of cleaved caspase-3 positive cells was calculated in ImageJ by counting cells with bright cleaved caspase-3 staining and dividing by the total number of DAPI-stained nuclei. Each bar comprises n = 153-553 cells pooled from multiple experiments.

**S6 Fig. RhoD-deficient cells display more frequent cytochrome *c* release, exhibit more caspase cleavage, and undergo more apoptosis following DNA damage**. **(A)** eHAP and RhoD^KO^ cells were treated with DMSO or 5μM etoposide for 6h and stained with Alexa488-AnnV (green), PI (red), and Hoescht (blue). Scale bar, 100μm. **(B)** Cells were treated with DMSO or etoposide and stained with caspase-3/7 green detection reagent to label cleaved DEVD (green) and Hoescht to stain DNA (blue). Scale bar, 100μm. **(C)** Cells were treated with DMSO or etoposide before being fixed and stained with a cytochrome *c* antibody (Cyto *c*; green), AIF antibody (Mito; red), and DAPI (DNA; blue). Scale bar, 25μm. **(D)** Cells were treated with DMSO for 6h or etoposide for 3 or 6h, and extracts were immunoblotted with anti-caspase-9 (Casp-9^Pro^ and Casp-9^Cleaved^) and anti-tubulin antibodies. For quantification, the caspase cleavage ratio was calculated by dividing the cleaved band intensity by the pro-caspase band intensity. Each point represents the mean ratio ±SD from 3 experiments. AU = Arbitrary Units. **(E)** Cells were treated with DMSO or etoposide for 6h, and extracts were immunoblotted with anti-cleaved caspase-3 (Casp-3^Cleaved^) and anti-tubulin antibodies. **(F)** Cells were treated with DMSO or etoposide for 6h before being fixed and stained with a cleaved caspase-3 antibody (Casp-3^Cleaved^; red), Alexa488-phalloidin (F-actin; green), and DAPI (DNA; blue). Scale bar, 25μm. The % of cleaved caspase-3 positive cells was calculated by counting cells with bright cleaved caspase-3 staining and dividing by the total number of DAPI-stained nuclei. Each bar comprises n = 117-410 cells pooled from multiple experiments. **p<0.01 (ANOVA, Tukey post-hoc tests).

**S1 Video. Live imaging of the apoptotic hallmarks of cell shrinkage, phosphatidylserine externalization, and membrane permeability after DNA damage**.

HAP1 cells were treated with 5μM etoposide for 4h and stained with Alexa488-AnnV (green), PI (red), and Hoescht (blue) prior to imaging at 37°C. Images were acquired every 30s for 2h and playback is at 10 frames/s. Scale bar, 25μm.

