## Supplemental Information for "The actin nucleation factors JMY and WHAMM enable a rapid p53-dependent pathway of apoptosis"

| S1 Table. Cell Lines | | | |
|---|---|---|---|
| ***Parental Cells*** | | | |
| Cell Line | Source | | |
| eHAP | Horizon Genomics (C669) | | |
| HAP1 | Horizon Genomics (C631) | | |
| HeLa | UC Berkeley Cell Culture Facility | | |
| U2OS | UC Berkeley Cell Culture Facility | | |
| ***HAP1 Derivatives*** | | | |
| KO Cell Line | Mutation | Predicted AAs | Source |
| Cortactin$^{KO}$ | 16bp deletion in exon 5 of 18 | 82/550, 49 post-shift | Horizon Genomics (HZGHC002628c003) |
| JMY$^{KO-1A}$ | 17bp deletion in exon 1 of 11 | 145/988, 4 post-shift | Horizon Genomics (HAP1_JMY_28380-03) |
| JMY$^{KO-1B}$ | 10bp deletion in exon 1 of 11 | 148/988, 30 post-shift | Horizon Genomics (HZGHC002630c002) |
| JMY$^{KO-2}$ | 2bp deletion in exon 2 of 11 | 362/988, 3 post-shift | Horizon Genomics (HZGHC002631c007) |
| N-WASP$^{KO}$ | 14bp deletion in exon 2 of 11 | 51/505, 12 post-shift | Horizon Genomics (HZGHC002632c003) |
| WASH Complex$^{KO}$ (CCDC53$^{KO}$) | 1bp insertion in exon 3 of 7 | 56/194, 6 post-shift | Horizon Genomics (HZGHC004026c001) |
| WAVE1$^{KO}$ | 11bp deletion in exon 5 of 11 | 83/559, 41 post-shift | Horizon Genomics (HZGHC0033300c010) |
| WAVE2$^{KO}$ | 46bp deletion in exon 3 of 9 | 64/498, 41 post-shift | Horizon Genomics (HZGHC003327c004) |
| WAVE3$^{KO}$ | 2bp deletion in exon 4 of 10 | 82/502, 35 post-shift | Horizon Genomics (HZGHC003325c004) |
| WAVE Complex$^{KO}$ (BRK1$^{KO}$) | 115bp insertion in exon 2 of 3 | 57/75, 45 post-shift | Horizon Genomics (HZGHC004027c003) |
| ***eHAP Derivatives*** | | | |
| KO Cell Line | Mutation | Predicted AAs | Source |
| RhoD$^{KO-2A}$ | 22bp deletion in exon 2 of 5 | 44/210, 8 post-shift | Horizon Genomics (HZGHC005085c003) |
| RhoD$^{KO-2B}$ | 1bp deletion in exon 2 of 5 | 48/210, 11 post-shift | Horizon Genomics (HZGHC005085c011) |
| WHAMM$^{KO-2}$ | 10bp deletion in exon 2 of 10 | 204/809, 35 post-shift | Horizon Genomics Mathiowetz et al., (2017) |
| WHAMM$^{KO-4}$ | 7bp deletion in exon 4 of 10 | 321/809, 17 post-shift | Horizon Genomics (HZGHC001060c001) |

**S1 Table. Cell Lines.**

| Gene | | Log2(FPKM $JMY^{KO-1A}$:HAP1) | | q-value |
|---|---|---|---|---|
| | | Mean | StDev | |
| JMY Interactors | EP300 | 0.17 | 0.02 | 0.999 |
| | STRAP | -0.31 | 0.13 | 0.999 |
| | TP53 | 0.57 | 0.08 | 0.999 |
| | MDM2 | -0.04 | 0.05 | 0.999 |
| Apoptotic Caspases | CASP2 | 0.09 | 0.14 | 0.999 |
| | CASP3 | -0.16 | 0.12 | 0.999 |
| | CASP6 | -0.31 | 0.01 | 0.999 |
| | CASP7 | -0.19 | 0.05 | 0.999 |
| | CASP8 | -0.04 | 0.14 | 0.999 |
| | CASP9 | -0.26 | 0.13 | 0.999 |
| Apoptosis Regulation | BIRC2 | -0.27 | 0.20 | 0.999 |
| | BIRC5 | 0.05 | 0.07 | 0.999 |
| | BIRC6 | 0.12 | 0.07 | 0.999 |
| | BCL2 | -0.39 | 0.33 | 0.999 |
| | BCL2L1 | 0.50 | 0.04 | 0.999 |
| | BCL2L10 | -0.04 | 0.25 | 0.999 |
| | BCL2L12 | 0.26 | 0.15 | 0.999 |
| | MCL1 | -0.33 | 0.04 | 0.999 |
| | BAX | -0.11 | 0.08 | 0.999 |
| | BAK1 | -0.06 | 0.13 | 0.999 |
| | BOK | 0.24 | 0.16 | 0.999 |
| | BID | -0.05 | 0.04 | 0.999 |
| | BCL2L11 | -0.20 | 0.17 | 0.999 |
| | BMF | 0.85 | 0.03 | 0.999 |
| | BAD | 0.00 | 0.48 | 0.999 |
| | BIK | -0.95 | 0.27 | 0.999 |
| | PMAIP1 | 0.09 | 0.14 | 0.999 |
| | BBC3 | 0.77 | 0.28 | 0.999 |
| | APAF1 | -0.14 | 0.04 | 0.999 |
| | CYCS | -0.17 | 0.26 | 0.999 |
| Cell Cycle Regulation | CDKN1A | 0.10 | 0.65 | 0.999 |
| | CDKN2A | 0.10 | 0.35 | 0.999 |
| Nucleation Promoting Factors | WASL | -0.50 | 0.08 | 0.999 |
| | WASH1 | 0.20 | 0.07 | 0.999 |
| | WASF1 | -0.43 | 0.06 | 0.999 |
| | WASF2 | 0.06 | 0.06 | 0.999 |
| | WASF3 | -0.13 | 0.20 | 0.999 |
| | WHAMM | -0.08 | 0.01 | 0.999 |
| | JMY | 0.05 | 0.29 | 0.999 |
| | NCKIPSD | -0.09 | 0.17 | 0.999 |
| | CTTN | 0.06 | 0.06 | 0.999 |
| Tandem Actin Monomer-binding Proteins of Nucleation | COBL | -0.03 | 0.08 | 0.999 |
| | SPIRE1 | 0.08 | 0.09 | 0.999 |
| | SPIRE2 | 0.77 | 0.35 | 0.999 |
| Formins | APC | -0.11 | 0.06 | 0.999 |
| | DIAPH1 | -0.18 | 0.02 | 0.999 |
| | DIAPH2 | 0.05 | 0.06 | 0.999 |
| | DIAPH3 | -0.07 | 0.13 | 0.999 |
| | DAAM1 | -0.29 | 0.14 | 0.999 |
| | FMN2 | 0.67 | 0.12 | 0.999 |
| | FMNL2 | -0.44 | 0.12 | 0.999 |
| | FMNL3 | 0.27 | 0.06 | 0.999 |
| | FHOD1 | 0.06 | 0.22 | 0.999 |
| | FHOD3 | 0.76 | 0.16 | 0.999 |
| | INF2 | 0.19 | 0.36 | 0.999 |
| Arp2/3 Complex | ACTR2 | -0.05 | 0.21 | 0.999 |
| | ACTR3 | 0.04 | 0.16 | 0.999 |
| | ARPC1A | -0.01 | 0.12 | 0.999 |
| | ARPC1B | -0.02 | 0.33 | 0.999 |
| | ARPC2 | -0.24 | 0.06 | 0.999 |
| | ARPC3 | -0.22 | 0.21 | 0.999 |
| | ARPC4 | -0.11 | 0.03 | 0.999 |
| | ARCP5 | -0.23 | 0.03 | 0.999 |
| | ARCP5L | -0.13 | 0.12 | 0.999 |

| Gene | | Log2 (FPKM $JMY^{KO-1A}$:HAP1) | | q-value |
|---|---|---|---|---|
| | | Mean | StDev | |
| Ras | DIRAS1 | 0.21 | 0.25 | 0.999 |
| | GEM | -0.31 | 0.55 | 0.999 |
| | HRAS | 0.13 | 0.12 | 0.999 |
| | KRAS | -0.08 | 0.10 | 0.999 |
| | MRAS | 0.69 | 0.09 | 0.999 |
| | NRAS | -0.21 | 0.12 | 0.999 |
| | NKIRAS1 | 0.00 | 0.24 | 0.999 |
| | NKIRAS2 | 0.07 | 0.13 | 0.999 |
| | RRAS | 0.35 | 0.10 | 0.999 |
| | RALA | -0.22 | 0.18 | 0.999 |
| | RALB | 0.09 | 0.16 | 0.999 |
| | RAP1A | -0.22 | 0.25 | 0.999 |
| | RAP1B | -0.08 | 0.21 | 0.999 |
| | RAP2A | -0.19 | 0.14 | 0.999 |
| | RAP2B | 0.06 | 0.23 | 0.999 |
| | RAP2C | -0.04 | 0.22 | 0.999 |
| | RASLL10B | 0.85 | 0.05 | 0.999 |
| | RASL11A | -0.41 | 0.18 | 0.999 |
| | RASL11B | -0.78 | 0.19 | 0.999 |
| | RHEB | -0.13 | 0.12 | 0.999 |
| | RHEBL1 | -0.14 | 0.36 | 0.999 |
| | RIT1 | -0.24 | 0.14 | 0.999 |
| | RRAS2 | 0.07 | 0.20 | 0.999 |
| Arf | ARF1 | 0.08 | 0.03 | 0.999 |
| | ARF3 | 0.20 | 0.09 | 0.999 |
| | ARF4 | 0.08 | 0.14 | 0.999 |
| | ARL4D | 0.24 | 0.37 | 0.999 |
| | ARF5 | 0.07 | 0.09 | 0.999 |
| | ARF6 | -0.14 | 0.05 | 0.999 |
| | ARFRP1 | 0.10 | 0.22 | 0.999 |
| | ARFRP2 | -0.74 | 0.20 | 0.999 |
| | ARL15 | -0.24 | 0.12 | 0.999 |
| | ARL2 | -0.27 | 0.12 | 0.999 |
| | ARL2BP | -0.03 | 0.11 | 0.999 |
| | ARL3 | 0.23 | 0.18 | 0.999 |
| | ARL4A | -0.15 | 0.14 | 0.999 |
| | ARL5A | -0.04 | 0.32 | 0.999 |
| | ARL6 | -0.22 | 0.07 | 0.999 |
| | ARL4C | 0.09 | 0.09 | 0.999 |
| | ARL8A | -0.21 | 0.21 | 0.999 |
| | SAR1A | -0.12 | 0.12 | 0.999 |
| | SAR1B | -0.05 | 0.18 | 0.999 |
| Rho | CDC42 | -0.01 | 0.14 | 0.999 |
| | RAC1 | -0.08 | 0.06 | 0.999 |
| | RAC2 | 1.67 | 0.77 | 0.999 |
| | RAC3 | -0.02 | 0.08 | 0.999 |
| | RHOA | -0.01 | 0.09 | 0.999 |
| | RHOB | -1.82 | 0.76 | 0.999 |
| | RHOBTB1 | -0.13 | 0.08 | 0.999 |
| | RHOBTB2 | -0.63 | 0.08 | 0.999 |
| | RHOBTB3 | -0.20 | 0.17 | 0.999 |
| | RHOC | 0.50 | 0.12 | 0.999 |
| | RHOD | 12.70 | 0.28 | 0.013 |
| | RHOF | 0.26 | 0.18 | 0.999 |
| | RHOG | 0.01 | 0.22 | 0.999 |
| | RHOQ | -0.17 | 0.15 | 0.999 |
| | RHOU | -0.62 | 0.20 | 0.999 |
| | RHOV | -0.12 | 0.13 | 0.999 |
| | RIF1 | -0.06 | 0.18 | 0.999 |
| | RND1 | 0.02 | 0.07 | 0.999 |
| | RND2 | 0.19 | 0.14 | 0.999 |
| | RAN | -0.10 | 0.08 | 0.999 |

| Gene | | Log2 (FPKM $JMY^{KO-1A}$:HAP1) | | q-value |
|---|---|---|---|---|
| | | Mean | StDev | |
| Rab | RAB1A | 0.05 | 0.12 | 0.999 |
| | RAB1B | -0.10 | 0.09 | 0.999 |
| | RAB2A | -0.34 | 0.11 | 0.999 |
| | RAB2B | 0.02 | 0.04 | 0.999 |
| | RAB3A | -0.25 | 0.42 | 0.999 |
| | RAB3B | -0.40 | 0.04 | 0.999 |
| | RAB3D | -0.20 | 0.13 | 0.999 |
| | RAB4A | -0.17 | 0.12 | 0.999 |
| | RAB4B | -0.49 | 0.47 | 0.999 |
| | RAB5A | -0.05 | 0.04 | 0.999 |
| | RAB5B | -0.12 | 0.03 | 0.999 |
| | RAB5C | 0.01 | 0.06 | 0.999 |
| | RAB6A | -0.11 | 0.02 | 0.999 |
| | RAB6B | -0.06 | 0.08 | 0.999 |
| | RAB7A | -0.30 | 0.08 | 0.999 |
| | RAB7L1 | -1.00 | 0.11 | 0.589 |
| | RAB8A | -0.07 | 0.09 | 0.999 |
| | RAB8B | 0.38 | 0.22 | 0.999 |
| | RAB9A | -0.39 | 0.22 | 0.999 |
| | RAB9B | -1.02 | 0.31 | 0.999 |
| | RAB10 | -0.17 | 0.12 | 0.999 |
| | RAB11A | -0.11 | 0.16 | 0.999 |
| | RAB11B | 0.18 | 0.29 | 0.999 |
| | RAB12 | -0.16 | 0.17 | 0.999 |
| | RAB13 | 0.06 | 0.18 | 0.999 |
| | RAB14 | -0.15 | 0.10 | 0.999 |
| | RAB15 | -0.88 | 0.14 | 0.648 |
| | RAB18 | 0.00 | 0.19 | 0.999 |
| | RAB20 | -0.59 | 0.34 | 0.999 |
| | RAB21 | 0.15 | 0.19 | 0.999 |
| | RAB22A | -0.48 | 0.10 | 0.999 |
| | RAB23 | -0.35 | 0.13 | 0.999 |
| | RAB24 | 0.05 | 0.19 | 0.999 |
| | RAB27A | -0.11 | 0.25 | 0.999 |
| | RAB28 | -0.07 | 0.16 | 0.999 |
| | RAB30 | 0.27 | 0.14 | 0.999 |
| | RAB31 | -0.06 | 0.05 | 0.999 |
| | RAB33A | -0.76 | 0.41 | 0.999 |
| | RAB33B | -0.02 | 0.06 | 0.999 |
| | RAB34 | 1.28 | 0.16 | 0.999 |
| | RAB35 | -0.04 | 0.08 | 0.999 |
| | RAB38 | -0.62 | 0.04 | 0.999 |
| | RAB39A | -0.18 | 0.79 | 0.999 |
| | RAB40A | 0.84 | 0.61 | 0.999 |
| | RAB40B | -0.25 | 0.07 | 0.999 |
| | RAB40C | 0.19 | 0.26 | 0.999 |
| | RAB42 | 0.17 | 0.22 | 0.999 |

| | Log2 (FPKM $JMY^{KO-1A}$:HAP1 | q-value |
|---|---|---|
| **Unchanged** | ----- | >0.05 |
| **Turned On** | >10 | <0.05 |
| **Up-regulated** | >1 | <0.05 |
| **Down-regulated** | <-1 | <0.05 |
| **Turned Off** | <-10 | <0.05 |

**S2 Table . Expression of genes encoding actin nucleation factors, cell cycle arrest proteins, canonical apoptosis regulators, and representative small G-proteins.**

**S3 Table. RNA and DNA Oligonucleotides**

***siRNAs***

| Target | Identifier |
| --- | --- |
| Control (Fig 3, 6, 9, S4) | Sigma SIC001 |
| JMY (Fig 3, S4) | A: SASI_HS01_00366091<br>B: SASI_HS01_00081989 |
| TP53 (Fig 6) | A: SASI_HS02_00302766<br>B: SASI_HS02_00302767<br>C: SASI_HS02_00302768 |
| RHOD (Fig 9) | A: SASI_HS01_00186024<br>B: SASI_HS01_00186023 |

***DNA Primers***

| Target | | Sequence |
| --- | --- | --- |
| β-ACTIN (Fig 8, 9) | F | GCTCGTCGTCGACAACGGCT |
| | R | GGTCATCTTCTCGCGGTTGG |
| CDKN1A (Fig 7) | F | GGCCCGTGAGCGATGGAACT |
| | R | GGAGTGGTAGAAATCTGTCATGCTGG |
| GAPDH (Fig 7, 9, 10) | F | CCTCCTGCACCACCAACTGC |
| | R | CTCCGACGCCTGCTTCACCA |
| RAB1A (Fig 8) | F | GTTATGCCAGTGAAAATGTCAACAAATTGT |
| | R | TGCTCTGAATTTTAACATTGGACTTCTCAG |
| RAB1B (Fig 8) | F | GCGAGAACGTCAATAAGCTCC |
| | R | GTCGATCTTGAGATTGGGCCGCT |
| RHOD (Fig 8, 9, 10) | F (nt 225-247) | AGATGACTATGACCGCCTGCGGC |
| | F (nt 124-147) | TTCCCCGAGAGCTACACCCCCACG |
| | R | ACGGCGTGGACGTTGTCATGGAG |
| WHAMM (Fig 8) | F | CTCCGTGCTCTGTCCTCATCCTCTCA |
| | R | CTAACCATCCCACTGGCCAGGGTCCT |

**S3 Table. RNA and DNA oligonucleotides.**

| **S4 Table. Immunofluorescence and Immunoblotting Reagents** | | | | |
|---|---|---|---|---|
| Target | Probe | | Conc. | Identifier |
| ***Primary Antibodies (Immunofluorescence)*** | | | | |
| Caspase-3$^{Cleaved}$ (Fig S5, S6) | anti-Cleaved Caspase-3 | Rabbit | 1:1,000 | Cell Signaling (9661) |
| Cytochrome C (Fig 5, 10, S6) | anti-Cytochrome C | Mouse | 1:500 | Cell Signaling (12963) |
| γH2AX (Fig S3) | anti-Phospho-Histone H2A.X | Rabbit | 1:1,000 | Cell Signaling (9718) |
| Mitochondria (Fig 5, 10, S6) | anti-AIF | Rabbit | 1:1,000 | Cell Signaling (5318) |
| p53 (Fig 6) | anti-p53 | Rabbit | 1:1,000 | Proteintech (10442-1-AP) |
| p21 (Fig 7) | anti-p21 | Rabbit | 1:250 | Cell Signaling (2947) |
| ***Primary Antibodies (Immunoblotting)*** | | | | |
| Casp-9$^{Pro}$ & Casp-9$^{Cleaved}$ (Fig 4, S6) | anti-Total Caspase-9 | Rabbit | 1:500 | Cell Signaling (9502) |
| Casp-3$^{Pro}$ & Casp-3$^{Cleaved}$ (Fig 4) | anti-Total Caspase-3 | Mouse | 1:500 | Cell Signaling (9662) |
| Casp-3$^{Cleaved}$ (Fig S6) | anti-Cleaved Caspase-3 | Rabbit | 1:250 | Cell Signaling (9661) |
| γH2AX (Fig S3) | anti-Phospho-Histone H2A.X | Rabbit | 1:1,000 | Cell Signaling (9718) |
| GAPDH (Fig 6) | anti-GAPDH | Mouse | 1:10,000 | Proteintech (60004-1-Ig) |
| JMY (Fig 2) | anti-JMY | Rabbit | 1:350 | Santa Cruz Biotech (sc-10027) |
| JMY (Fig 3, S4) | anti-JMY | Rabbit | 1:1,000 | Proteintech (25098-1-AP) |
| p53 (Fig 6) | anti-p53 | Mouse | 1:250 | Cell Signaling (9282) |
| p53$^{phospho-S15}$ (Fig 6) | anti-Phospho-p53 (S15) | Rabbit | 1:500 | Cell Signaling (9284) |
| p53$^{acetyl-K382}$ (Fig 6) | anti-Acetyl-p53 (K382) | Rabbit | 1:500 | Cell Signaling (2525) |
| Tubulin (Fig 2, 3, 4, 6, 10, S1, S4, S6) | anti-Beta-Tubulin | Mouse | 1:10,000 | Developmental Studies Hybridoma Bank (E7) |
| WHAMM (Fig 2) | anti-WHAMM | Rabbit | 1:1,000 | Shen et al., (2012) |
| ***Secondary Antibodies (Immunofluorescence)*** | | | | |
| Mouse IgG | Alexa 488, 555 anti-mouse | Goat | 4 μg/ml | Life Technologies (e.g. A21424) |
| Rabbit IgG | Alexa 488, 555 anti-rabbit | Goat | 4 μg/ml | Life Technologies (e.g. A11034) |
| ***Secondary Antibodies (Immunoblotting)*** | | | | |
| Mouse IgG | HRP anti-Mouse | Sheep | 1:10,000 | GE Healthcare (NXA931) |
| Rabbit IgG | HRP anti-Rabbit | Donkey | 1:10,000 | GE Healthcare (NA934V) |
| Mouse IgG | IRDye 680, 800 anti-Mouse | Donkey | 0.05 μg/ml | LI-COR (e.g. 926-32212) |
| Rabbit IgG | IRDye 680, 800 anti-Rabbit | Donkey | 0.05 μg/ml | LI-COR (e.g. 926-32213) |
| ***Molecular Probes (Fluorescence)*** | | | | |
| Target | Probe | | Conc. | Identifier |
| Active Caspase-3/7 (Fig 4, 10, S6) | CellEvent Caspase-3/7 Green Detection Reagent | | 5 μM | Invitrogen (C10423) |
| DNA (Fig 1, 2, 3, 4, 6, 7, 9, 10, S1, S4, S6) | Hoescht 33342 Solution | | 2 μg/ml | Thermo Scientific (62249) |
| DNA (Fig 5, 6, 7, 10, S3, S5, S6) | 4′,6-diamidino-2-phenylindole (DAPI) | | 1 μg/ml | Invitrogen (D1306) |
| F-actin (Fig S3, S5, S6) | Alexa488-Phalloidin | | 0.2 U/ml | Invitrogen (A12379) |
| Nucleic Acids (Fig 1, 2, 3, 6, 7, 9, 10, S1, S4, S6) | Propidium Iodide | | 2 μg/ml | Invitrogen (P3566) |
| Phosphatidylserine (Fig 1, 2, 3, 6, 7, 9, 10, S1, S4, S6) | Alexa488-Annexin V | | 8 μM | Invitrogen (A13201) |

**S4 Table. Immunofluorescence and immunoblotting reagents.**

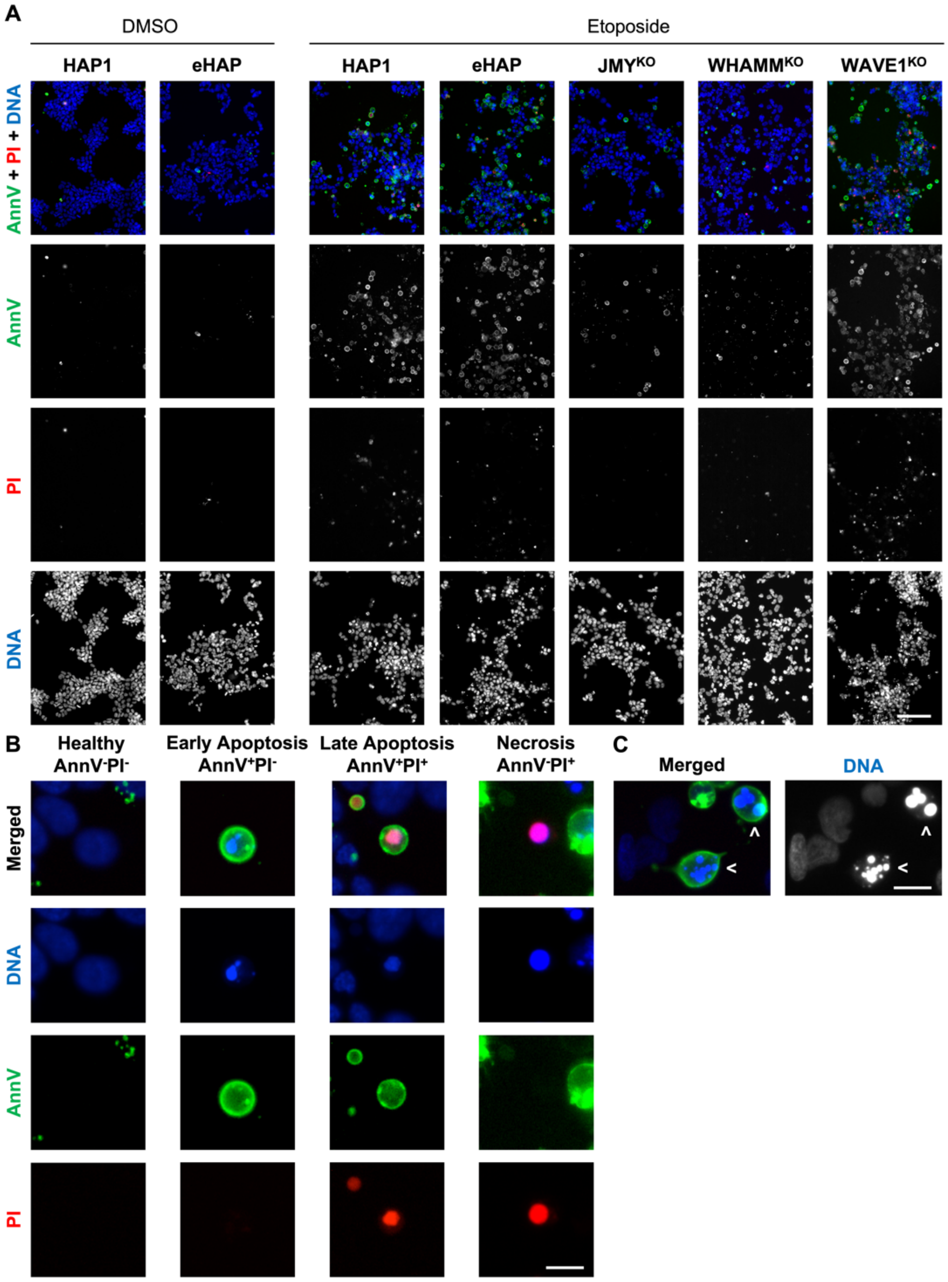


**S1 Fig. Cells lacking the WASP-family members JMY or WHAMM undergo less apoptosis.**

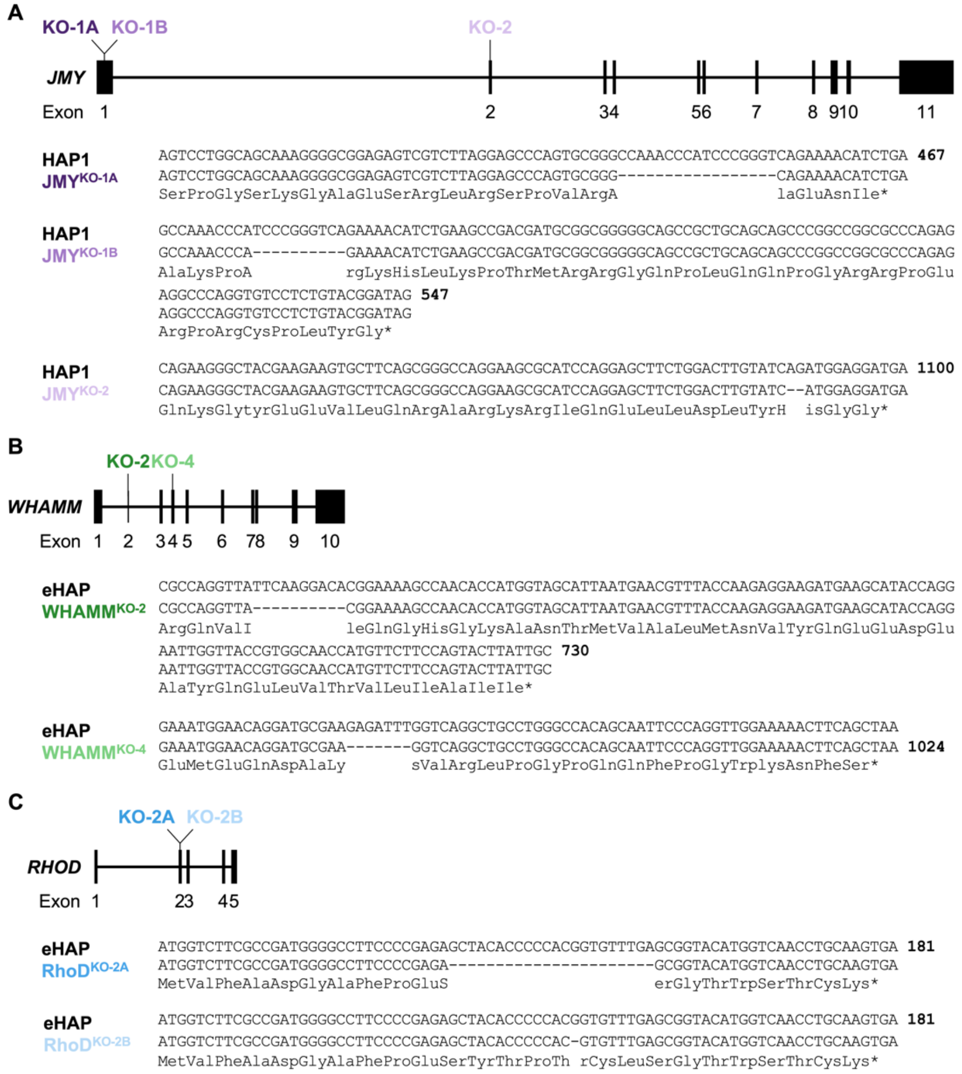


**S2 Fig. JMY, WHAMM, and RhoD knockout cell lines contain loss-of-function mutations derived from frameshifts or altered splicing.**

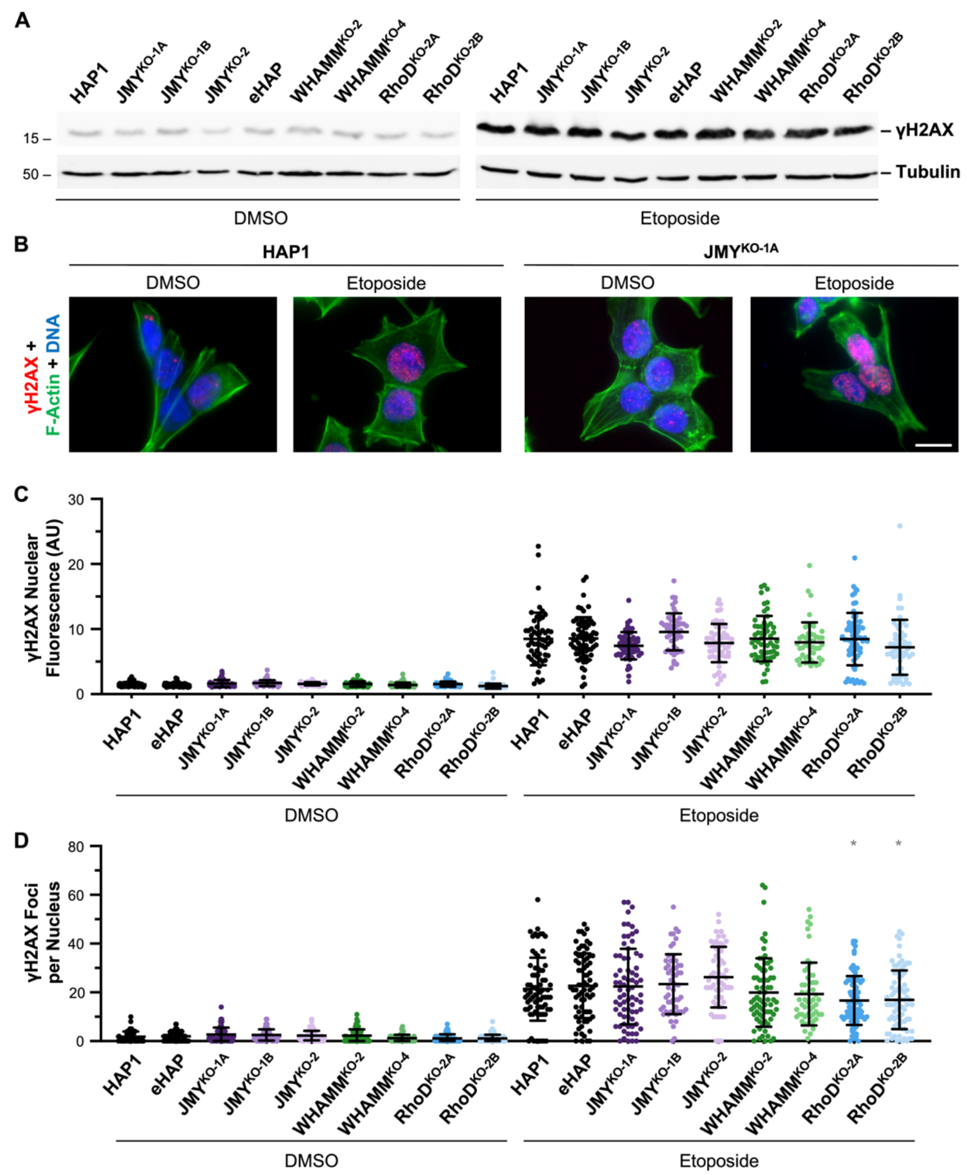


**S3 Fig. Parental and knockout cell lines exhibit high levels of γH2AX expression and clustering in response to DNA damage.**

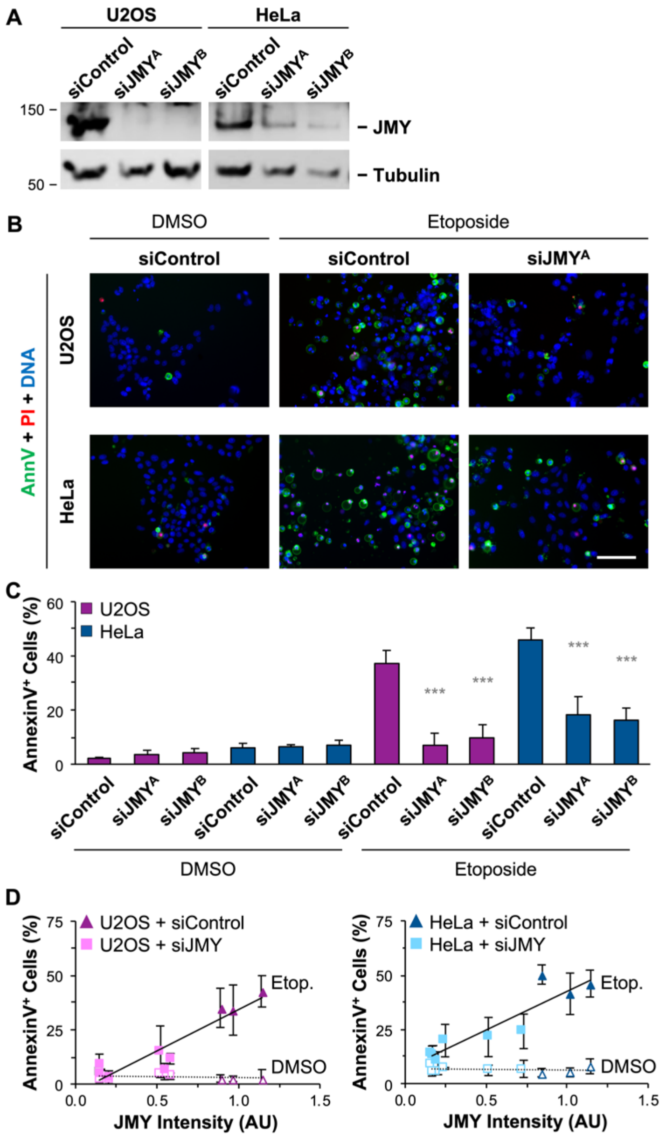


**S4 Fig. Transient JMY depletion in multiple cell lines results in less apoptosis following DNA damage.**

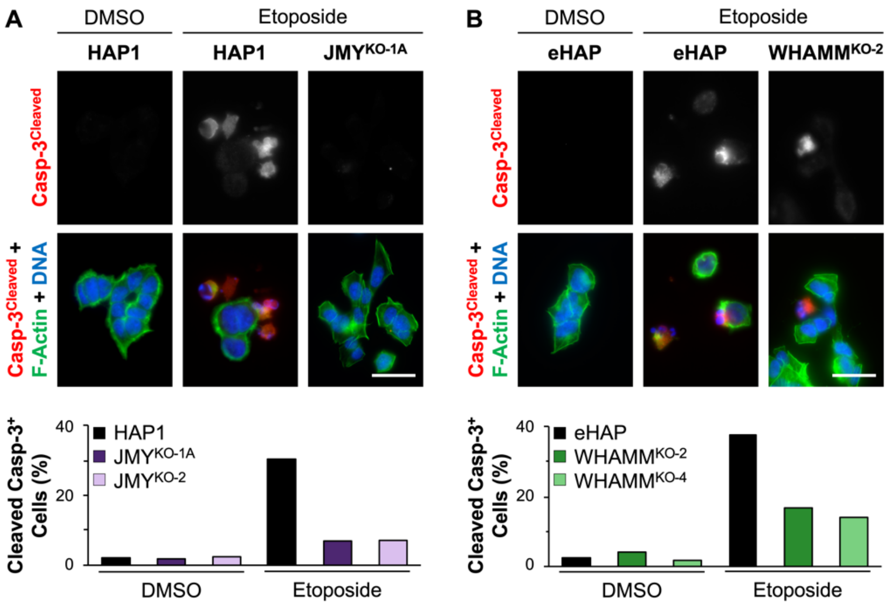


**S5 Fig. Caspase-3 cleavage is inefficient in JMY- and WHAMM-knockout cells.**

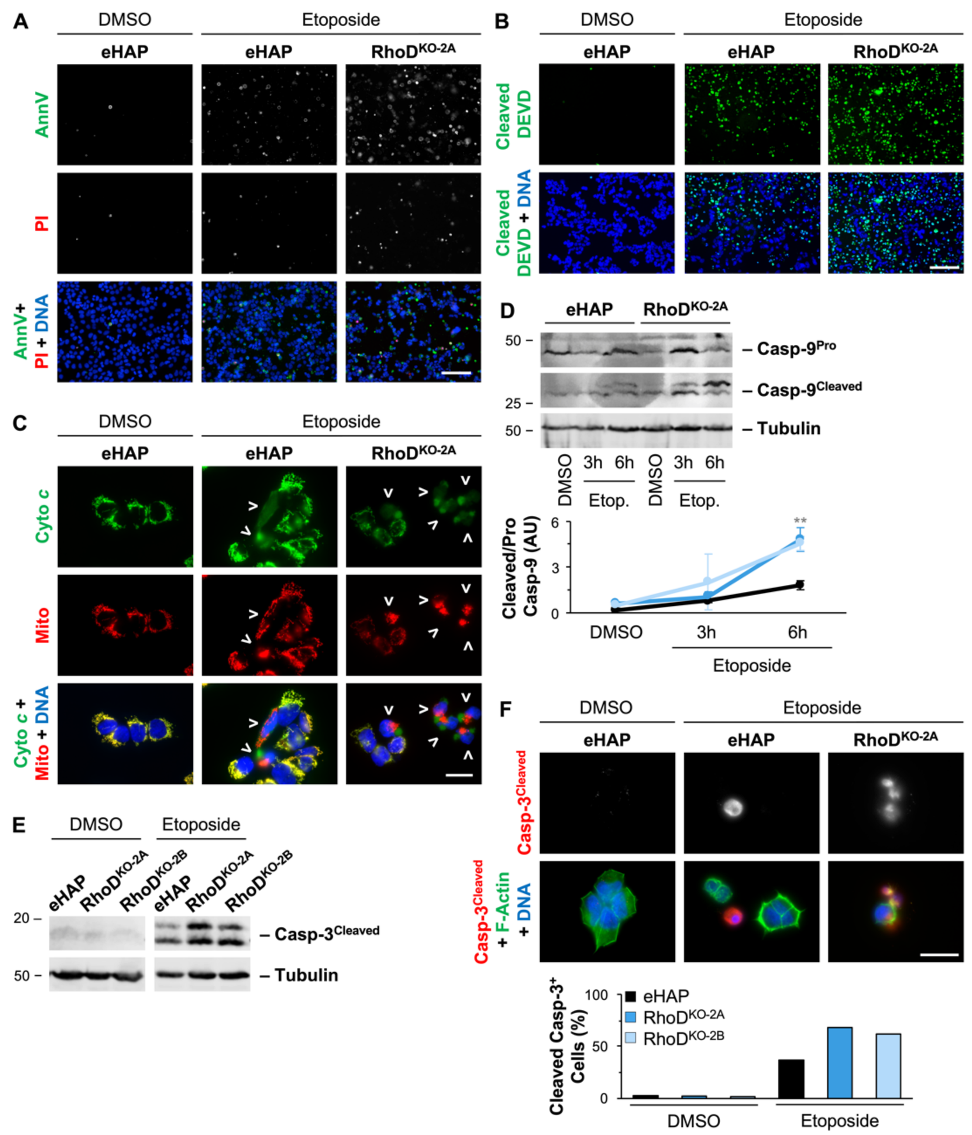


**S6 Fig. RhoD-deficient cells display more frequent cytochrome *c* release, exhibit more caspase cleavage, and undergo more apoptosis following DNA damage.**
